# The functional relevance of task-state functional connectivity

**DOI:** 10.1101/2020.07.06.187245

**Authors:** Michael W. Cole, Takuya Ito, Carrisa Cocuzza, Ruben Sanchez-Romero

## Abstract

Resting-state functional connectivity has provided substantial insight into intrinsic brain network organization, yet the functional importance of task-related change from that intrinsic network organization remains unclear. Indeed, such task-related changes are known to be small, suggesting they may have only minimal functional relevance. Alternatively, despite their small amplitude, these task-related changes may be essential for the human brain’s ability to adaptively alter its functionality via rapid changes in inter-regional relationships. We utilized activity flow mapping – an approach for building empirically-derived network models – to quantify the functional importance of task-state functional connectivity (above and beyond resting-state functional connectivity) in shaping cognitive task activations in the (female and male) human brain. We found that task-state functional connectivity could be used to better predict independent fMRI activations across all 24 task conditions and all 360 cortical regions tested. Further, we found that prediction accuracy was strongly driven by individual-specific functional connectivity patterns, while functional connectivity patterns from other tasks (task-general functional connectivity) still improved predictions beyond resting-state functional connectivity. Additionally, since activity flow models simulate how task-evoked activations (which underlie behavior) are generated, these results may provide mechanistic insight into why prior studies found correlations between task-state functional connectivity and individual differences in behavior. These findings suggest that task-related changes to functional connections play an important role in dynamically reshaping brain network organization, shifting the flow of neural activity during task performance.

**Significance Statement:** Human cognition is highly dynamic, yet the human brain’s functional network organization is highly similar across rest and task states. We hypothesized that, despite this overall network stability, task-related changes from the brain’s intrinsic (resting-state) network organization strongly contribute to brain activations during cognitive task performance. Given that cognitive task activations emerge through network interactions, we leveraged connectivity-based models to predict independent cognitive task activations using resting-state versus task-state functional connectivity. This revealed that task-related changes in functional network organization increased prediction accuracy of cognitive task activations substantially, demonstrating their likely functional relevance for dynamic cognitive processes despite the small size of these task-related network changes.

## Introduction

We and others recently found using functional MRI (fMRI) that the statistical relationship between brain regions – functional connectivity (FC) – is similar between resting state and a variety of task states (Cole et al., 2014; Krienen et al., 2014; Gratton et al., 2018). Despite the small task-related changes in FC some statistically reliable changes between rest and task states have been observed (Cole et al., 2014; Krienen et al., 2014; Gratton et al., 2018; Ito et al., 2020a). The functional relevance of these relatively small task-related FC changes remains unclear, despite the clear relationship between these FC changes and the transition from rest to task states.

We recently developed an approach to quantify the functional relevance of resting-state FC via incorporating FC into simple network models that predict task-evoked activations (Cole et al., 2016). Here we sought to extend this approach to task fMRI data to assess the functional relevance of task-state FC. The core mechanisms underlying these models are the same as those used in most biological and artificial neural network models: the propagation and activation rules (**Figure 1A**) (McClelland and Rogers, 2003; Ito et al., 2020b). The propagation rule specifies that a distal node’s activity influences a given target node via a connection weight, while the activation rule specifies that the incoming activity will be summed prior to passing through a function (typically a nonlinearity) to determine the output activity of the target node. The activity flow mapping approach (**Figure 1B**) builds on this framework, incorporating empirical FC and activation estimates to build a predictive model for each to-be-predicted region (one at a time). Since the to-be-predicted region’s activity level was empirically observed we can then test the accuracy of our model by comparison to empirical data. Thus, by predicting a variety of cognitive task-evoked activations (e.g., across regions and across task conditions), activity flow mapping provides a means to empirically test the functional (cognitive and computational) relevance of the connectivity estimates included in activity flow models (Ito et al., 2020b).

**Figure 1.**
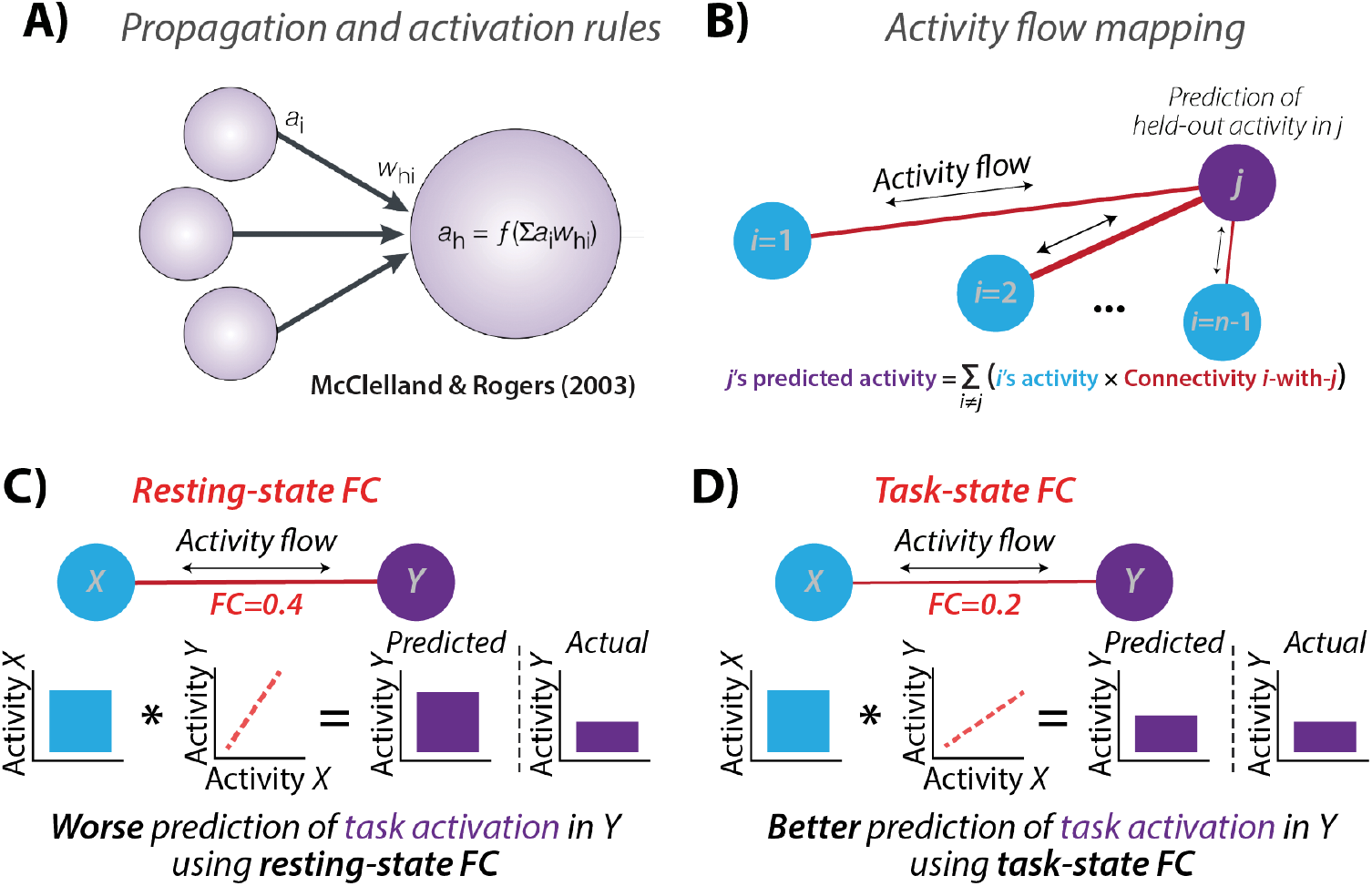
Predicted boost to activity flow-based predictions using task-state functional connectivity (FC). **A**) The propagation and activation rules used in neural network modeling provide a framework for modeling the flow of neural activity through networks. The propagation rule can be visualized via the arrows connecting distal nodes (such as region *i*) to a given target node *h* via a connection with strength *w_hi_*. The activation rule can be visualized via the summing of incoming activity in the target node *h*, then passing through an activation function *f*. Equation for computing the activity level *a* in node *h*: *a_h_* = *f*(*Σa_i_w_hi_*). Adapted from McClelland & Rogers (2003). **B**) We recently developed the activity flow mapping framework, applying neural network modeling to empirical connectivity (and activity) estimates. We showed that activity flow mapping can predict independent (held-out) task activations using resting-state FC (Cole et al., 2016; Ito et al., 2017). Figure from Cole et al. (2016). **C**) An illustration of simplified activity flow prediction of task activity in neural population *Y* based on task activity in neural population *X*, based on the resting-state FC between *X* and *Y*. **D**) The hypothesized boost in prediction accuracy by using the FC estimates from the same state as the task activity estimates. Note that the to-be-predicted task activity levels are carefully removed prior to estimating task-state FC (Cole et al., 2019) to avoid circularity.

Here we utilized activity flow mapping to test the functional relevance of task-state FC, especially as task-state FC differs from resting-state FC. The critical test was whether task-state FC increased activity-flow-based prediction accuracy across a variety of task conditions. Such an observation would be non-trivial for several reasons. First, the relatively small changes between task-state FC and resting-state FC makes it unclear whether task-state FC would produce significantly better task-evoked activation prediction accuracies. Second, we recently validated an improved method for the subtraction of mean task-evoked activations prior to task-state FC estimation with fMRI data (Cole et al., 2019), which is the same approach used in the noise correlation literature to characterize the functional relationship between neurons and neural populations (typically used in non-human animal neuroscience) (Ito et al., 2020a). This better ensures, relative to many prior studies, that the to-be-predicted activations have no direct effect on FC estimates (reducing potential circularity) (Cole et al., 2019).

Third, we recently replicated the non-human animal literature in finding that most human fMRI FC estimates decreased from rest to task (Ito et al., 2020a). This appears to run counter to the common intuition that FC should increase when two neural populations interact. One possibility is that – rather than FC increasing – the activity flow between two neural populations increases as the linear relationship between the populations (i.e., their FC) decreases (**Figure 1C & 1D**). This would be consistent with the well-supported possibility that the relationship between neural populations is a sigmoidal transfer function (Wilson and Cowan, 1972; Hopfield, 1982; Ito et al., 2020a). We recently used spiking and neural mass models to show that a sigmoidal transfer function between brain regions – which quenches variance at high and low activity levels – was enough to account for the observed FC decreases with increasing task-evoked neural activity (Ito et al., 2020a) (**Figure 2**). However, regardless of whether the sigmoidal transfer function is the mechanism underlying the relationship between neural populations, observing improved prediction accuracies using task-state FC would provide evidence that the observed changes in FC from rest to task are likely functionally relevant (rather than statistical artifacts). This is due to the clear functional relevance of what is being predicted: task-evoked activations across a variety of brain regions and task conditions (e.g., visual cortex responses in visual tasks, motor cortex responses in motor tasks). Indeed, task activations (neural activity level changes) can cause perceptual, motor, and cognitive processes (e.g., due to neural stimulation) (Valero-Cabré et al., 2017), and likely depend on state-dependent network reconfigurations.

**Figure 2.**
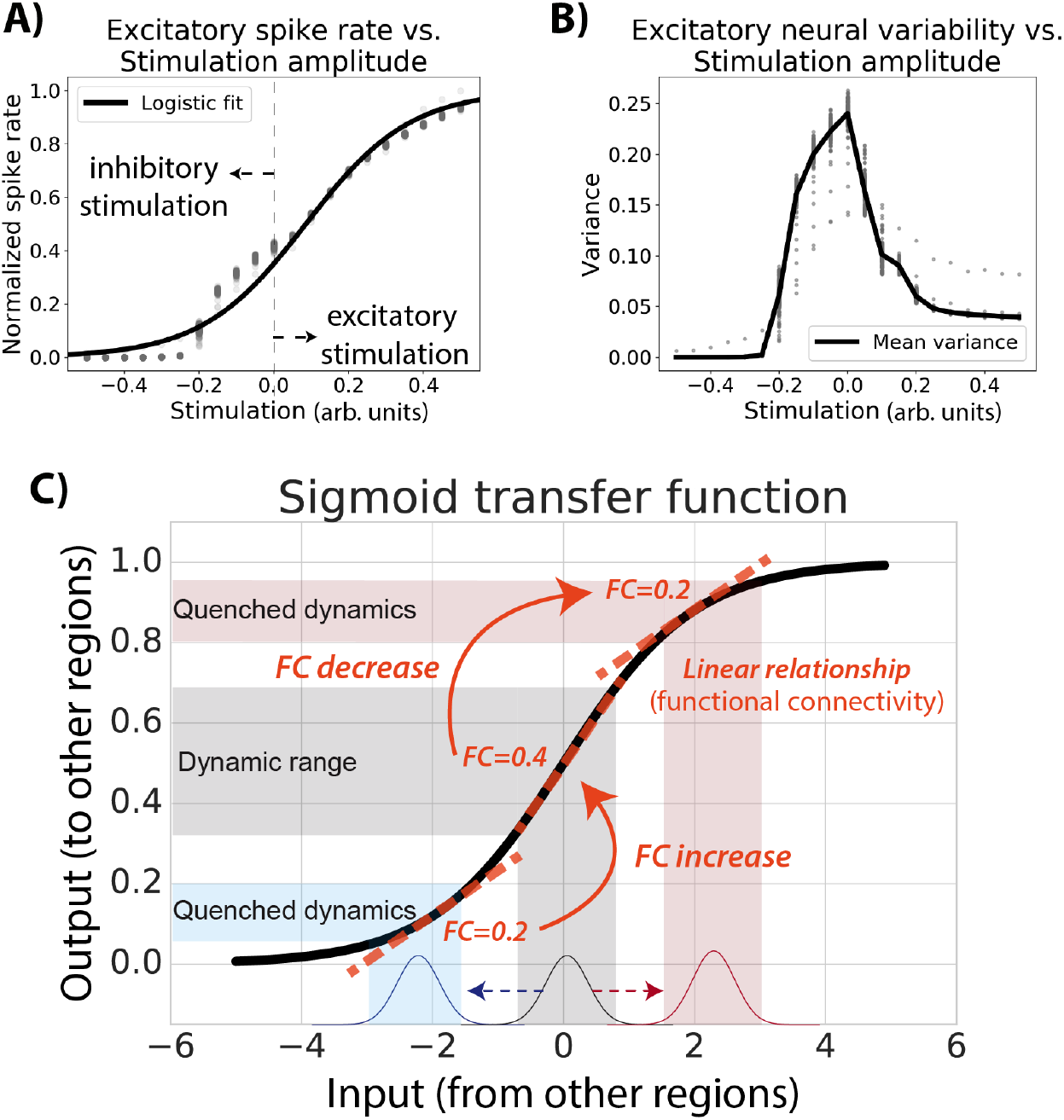
Neural populations are thought to interact via sigmoidal activation functions, helping explain widespread FC decreases from resting to task states. **A**) Spiking model simulation results from Ito et al. (2020a), showing sigmoidal relationship between inputs and outputs of neural populations. **B**) Activity increases and decreases from resting state in the spiking model (Ito et al., 2020a) resulted in decreased variance and correlations in simulated excitatory neurons. Empirical results in both spiking populations and fMRI show that variance and correlations decrease as activity levels increase or decrease from resting state (He, 2013; 2020a). **C**) There is substantial evidence that biological neural populations have a sigmoidal relationship, which could help explain well-known neural variability quenching effects from rest to task (He, 2013) as well as reductions in FC from rest to task (Ito et al., 2020a). The relationship between activity levels across neural populations (i.e., FC) changes as a function of overall activity levels.

The critical test of our hypothesis that task-state FC is functionally relevant was whether task-evoked activations are better predicted by network models parameterized by task-state FC than those parameterized by resting-state FC. This test would demonstrate that task-related network reconfigurations facilitate the propagation of task-related activations, which are commonly thought of as the primary neural substrate of cognitive processes (Varoquaux et al., 2018). We carried out a variety of tests of this hypothesis using fMRI data from the Human Connectome Project, which provided a large number of task conditions (24) for activity flow models to predict, as well as a large amount of task-state and resting-state data per subject for estimating FC. We began by testing for task-state vs. resting-state FC using the field-standard Pearson correlation FC measure. We then tested whether better accounting for causal confounds in activity flow models improved prediction accuracy. Finally, we tested various factors contributing to task-state FC prediction accuracy, such as testing whether task-state FC from other tasks (task-general FC) could improve predictions as well. Confirmation that task-state FC consistently improves task activation prediction across a variety of task conditions would suggest an important role for task-driven network changes in producing the task activations underlying perceptual, motor, and cognitive processes.

## Materials and Methods

### Activity flow mapping

We previously developed and validated activity flow mapping as a method to help determine the mechanistic role of empirically estimated connectivity in producing neural activations (**Figure 1B**) (Cole et al., 2016). Activity flow mapping involves three steps. First, a network model is constructed for each node using empirical connectivity estimates. Second, activity flows are simulated over each model’s connections to produce predicted activity in each node. Third, prediction accuracy is assessed by comparing the predicted activity to the actual empirically observed activity for each node. Prediction accuracy quantifies the likely validity of each activity flow model. All activity flow mapping analyses used the publicly available Brain Activity Flow (“Actflow”) Toolbox (https://colelab.github.io/ActflowToolbox/), version 0.2.5.

Based on neural network simulations (**Figure 1A**), the activity flow simulation step involves estimating net input to each target region by multiplying each other brain region’s task-related activation amplitude (analogous to the amount of neural activity) by its FC with the target region (analogous to aggregate synaptic strength):

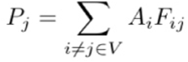

where *P* is the predicted mean activation for region *j* in a given task, *A_i_* is the actual mean activation for region *i* in a given task (a beta value estimated using a general linear model), *i* indexes all brain regions (vector *V*) with the exception of region *j*, and *F_ij_* is the FC estimate between region *i* and region *j*. This algorithm results in a matrix with predicted activations across all nodes and task conditions. Prediction accuracy was assessed by comparing predicted to actual empirical activation patterns using multiple approaches (**Table 1**). Unless specified otherwise, comparisons were made across both nodes and conditions simultaneously. This was accomplished by collapsing each of the predicted and actual node-by-condition matrices into a single vector of numbers prior to comparison.

Activity flow mapping was developed in accordance with several principles that facilitate its utility for scientific inferences. First, the approach is agnostic to the particular form of connectivity, so it can be used with any form of FC, effective connectivity, or structural connectivity (i.e., any estimate of the routes of activity flow between nodes). This provides the approach with extensive flexibility. Second, activity flow mapping can be seen as a method to test the validity of connectivity approaches in that it tests for model accuracy via prediction of independent data. Third, unlike standard connectivity benchmarks that depend on test-retest reliability, activity flow mapping assesses the mechanistic role of network models in producing neural activations. Thus, model evaluation is not performed on data of the same form (e.g., connectivity estimates) but rather of a mechanistically distinct form: neural activations. Thus, activity flow mapping can be seen as potentially providing insights into computation and the emergence of cognitive information (represented via activation levels/patterns) in each node (Ito et al., 2020b).

**Table 1.** Overview of predicted-to-actual assessment measures.

| Predicted-to-actual measure | Description |
| --- | --- |
| Pearson correlation (r) | Scaled similarity of a predicted task activity pattern to an actual activity pattern. Unaffected by the prediction being off by a multiplicative factor (e.g., r would be high even if a prediction is 2 times higher than actual but with a similar pattern). |
| R<sup>2</sup> (coeff. of determination) | Unscaled similarity of a predicted task activity pattern to an actual activity pattern (not the same as r<sup>2</sup>). When multiplied by 100, percentage of the to-be-predicted data's unscaled variance, ranging from negative infinity (because prediction errors can be arbitrarily large) to positive 1. An R<sup>2</sup> of 0 is equivalent to accurately predicting the mean of the data only. This is a standard definition of R<sup>2</sup> in machine learning for quantifying continuous predictions. |
| Node-wise comparison | Similarity of a predicted spatial activity pattern to an actual spatial activity pattern based on node-to-node variance. This can be computed separately for each task state of interest, but collapses across spatial locations. This was used to ensure replication of results across task states. |
| Condition-wise comparison | Similarity of a predicted activity pattern to an actual activity pattern based on condition-to-condition variance. This can be computed separately for each node, but collapses across task conditions. This can be thought of as a node's response profile or population receptive field. |
| Compare-then-average | Predicted-to-actual similarity computed for each individual subject separately, prior to averaging similarity estimates across individuals for a group result. This better characterizes the true predicted-to-actual similarity, since averaging across subjects blurs features of the data due to individual differences in anatomy (among other features). **Unless otherwise noted, all analyses used the compare-then-average approach.** |
| Average-then-compare | Predicted-to-actual similarity computed after averaging predicted activations across all subjects and actual activations across all subjects. While this likely distorts the true predicted-to-actual similarity to some extent, it has the advantage of better signal-to-noise due to averaging of much more data (176 times the data of compare-then-average for this study). It is also more difficult to compute valid p-values with this approach, since (unlike compare-then-average) inter-subject variance cannot be taken into account to help ensure results will generalize to new subjects. |

### Experimental Design and Statistical Analyses

See subsection “Data Collection” below for details on the experimental design. Paired t-tests (paired by subject) were used for statistical tests when possible. See subsections “FC estimation”, “Task activation level estimation", and “Family-wise error correction for multiple comparisons” for further details on statistical analyses.

### Data collection

We used the Washington University-Minnesota Consortium Human Connectome Project (HCP) young adult publicly available dataset (Van Essen et al., 2013) (available at https://www.humanconnectome.org/study/hcp-young-adult). Participants were recruited from Washington University (St. Louis, MO) and the surrounding area. All participants gave informed consent. We selected 352 low-motion subjects (with no family relations) from the “1200 Subjects” HCP release. We split the 352 subjects into two cohorts of 176 subjects: an exploratory cohort (99 females) and a replication cohort (84 females). The exploratory cohort had a mean age of 29 years of age (range=22-36 years of age), and the replication cohort had a mean age of 28 years of age (range=22-36 years of age). These 352 participants were selected by excluding those with any fMRI run in which more than 50% of TRs had greater than 0.25mm framewise displacement, or if a family relation was already included (or if they had no genotyping to verify family relation status). A full list of the 352 participants used in this study will be included as part of the code release.

Whole-brain echo-planar imaging acquisitions were acquired with a 32 channel head coil on a modified 3T Siemens Skyra MRI with TR = 720 ms, TE = 33.1 ms, flip angle = 52°, BW = 2290 Hz/Px, in-plane FOV = 208 × 180 mm, 72 slices, 2.0 mm isotropic voxels, with a multi-band acceleration factor of 8 (Sotiropoulos et al., 2013). Data were collected over two days. On each day 28 minutes of rest (eyes open with fixation) fMRI data across two runs were collected (56 minutes total), followed by 30 minutes of task fMRI data collection (60 minutes total). Resting-state data collection details for this dataset can be found elsewhere (Smith et al., 2013), as can task data details (Barch et al., 2013).

Each of the seven tasks was collected over two consecutive fMRI runs. The seven tasks consisted of an emotion cognition task, a gambling reward task, a language task, a motor task, a relational reasoning task, a social cognition task, and a working memory task. Briefly, the emotion cognition task required making valence judgements on negative (fearful and angry) and neutral faces. The gambling reward task consisted of a card guessing game, where subjects were asked to guess the number on the card to win or lose money. The language processing task consisted of interleaving a language condition, which involved answering questions related to a story presented aurally, and a math condition, which involved basic arithmetic questions presented aurally. The motor task involved asking subjects to either tap their left/right fingers, squeeze their left/right toes, or move their tongue. The reasoning task involved asking subjects to determine whether two sets of objects differed from each other in the same dimension (e.g., shape or texture). The social cognition task was a theory of mind task, where objects (squares, circles, triangles) interacted with each other in a video clip, and subjects were subsequently asked whether the objects interacted in a social manner. Lastly, the working memory task was a variant of the N-back task. Further details on the HCP task paradigms can be found elsewhere (Barch et al., 2013).

### Data preprocessing

Preprocessing was carried out identically to another recent study that used HCP data (Ito et al., 2020a). Minimally preprocessed data for both resting-state and task fMRI were obtained from the publicly available HCP data. Minimally preprocessed surface data was then parcellated into 360 brain regions using the Glasser (2016) atlas. We performed additional standard preprocessing steps on the parcellated resting-state fMRI and task-state fMRI data. This included removing the first five frames of each run, de-meaning and de-trending the time series, and performing nuisance regression on the minimally preprocessed data. Nuisance regression was based on empirical validation tests by Ciric et al. (2017) to reduce effects of motion and physiological noise. Specifically, six primary motion parameters were removed, along with their derivatives, and the quadratics of all regressors (24 motion regressors in total). Physiological noise was modeled using aCompCor on time series extracted from the white matter and ventricles (Behzadi et al., 2007). For aCompCor, the first 5 principal components from the white matter and ventricles were extracted separately and included in the nuisance regression. In addition, we included the derivatives of each of those components, and the quadratics of all physiological noise regressors (40 physiological noise regressors in total). The nuisance regression model contained a total of 64 nuisance parameters. Note that aCompCor was used in place of global signal regression, given evidence that it has similar benefits as global signal regression for removing artifacts (Power et al., 2018) but without regressing gray matter signals (mixed with other gray matter signals) from themselves, which may result in false correlations (Murphy et al., 2009; Power et al., 2017).

Task data for task-state FC analyses were additionally preprocessed using a general linear model (GLM). The mean evoked task-related activity for each of the 24 task conditions was removed by fitting the task timing (block design) for each condition. This was accomplished using the exact same canonical hemodynamic response regressors as used for the task activation estimates, fit simultaneously with a finite impulse response (FIR) model (Cole et al., 2019). This was critical for removing data analysis circularity (Kriegeskorte et al., 2009), separating the to-be-predicted task activations from the task-state FC estimates used to predict them (**Figure 3**). Including the exact same regressors as used for task activation estimation was important to ensure statistical non-circularity in the activity flow mapping analyses. We used a FIR regressors in addition to the canonical hemodynamic response regressors given recent evidence suggesting that the FIR model reduces both false positives and false negatives in the identification of FC estimates (Cole et al., 2019). Further, FIR models are more flexible, removing cross-event/block variance much better than alternative approaches to better reduce the chance of analysis circularity. Each set of FIR regressors included a lag extending 25 time points after task block offset to account for the post-event hemodynamic undershoot.

**Figure 3.**
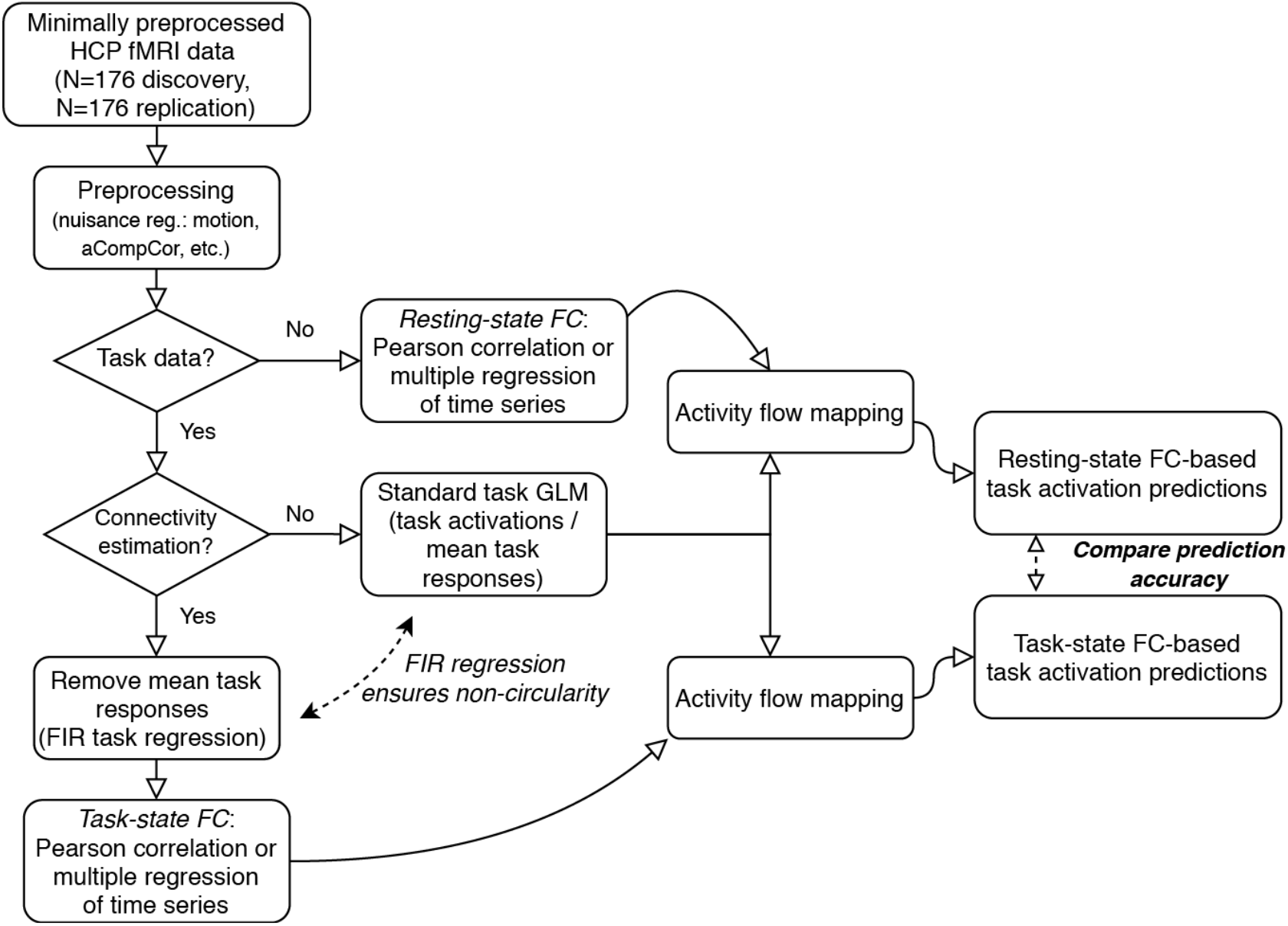
The fMRI data processing workflow for comparing task activation predictions based on task-state vs. resting-state FC. Publically available Human Connectome Project fMRI data (young adult dataset) (N=352) was split into separate discovery and replication datasets (N=176 each). Circularity is carefully avoided in the predictions by 1) finite impulse response (FIR) regression to remove cross-block mean task responses prior to task-state FC estimation, and 2) removal of each to-be-predicted brain region from the set of predictors in the activity flow mapping step. The goal of the primary analyses is to compare task activation predictions based on resting-state FC to predictions based on task-state FC.

### FC estimation

All FC estimates were computed using the publicly available Brain Activity Flow (“Actflow”) Toolbox (https://colelab.github.io/ActflowToolbox/). The initial analyses estimated FC using Pearson correlations between time series (averaging across voxels within each region) from all pairs of brain regions. All computations involving Pearson correlations used Fisher’s z-transformed values, which were reconverted to r-values for reporting purposes.

We used multiple linear regression (the LinearRegression function in the Scikit-learn Python package) as an alternative to Pearson correlation. This involved computing a linear model for each to-be-predicted region separately. Time series from all other regions were used as predictors of the to-be-predicted region’s time series. The resulting betas – which were directional from the predictor regions to the predicted region – were then used as FC estimates in the activity flow mapping algorithm. Note that beta estimate directionality reflects optimal linear scaling of the source time series to best match the target time series, not necessarily the direction of activity flow. Regularized regression was used when there were fewer time points than nodes. Specifically, principal component regression was used, including the maximum number of components possible (e.g., 199 components if 200 time points were available).

The number of time points contributing to each FC estimate was matched for all analyses unless otherwise specified. This typically involved restricting the amount of resting-state data used for estimating resting-state FC based on the limited amount of task-state data for the to-be-compared task condition. This ensured the task-state FC and resting-state FC were equated in terms of the amount of data contributing to their estimates, increasing the validity of comparisons between the two types of FC estimates.

The Cole-Anticevic Brain-wide Network Partition (CAB-NP) was used for visualization of network structure (Ji et al., 2019) (available at https://github.com/ColeLab/ColeAnticevicNetPartition).

### Task activation level estimation

Task-evoked activation amplitudes were estimated using a standard general linear model. The SPM software canonical hemodynamic response function was used for general linear model estimation, given that all tasks involved block designs.

### Family-wise error correction for multiple comparisons

We used the MaxT non-parametric permutation testing approach to correct for multiple comparisons (Nichols and Holmes, 2002). This involved 1000 permutations to create a null distribution of t-values, which the actual (paired t-test) t-values could be compared to compute a non-parametric p-value that was family-wise error corrected for multiple comparisons.

### Assessing individual-specific FC changes

Individual-specific and state-specific FC effects were evaluated by predicting task-evoked activations during only the second run of each task. This allowed us to distinguish between the effects of an individual’s task-state FC generally (estimated from the first run of a given task) versus an individual’s task-state FC from the same run as the task-evoked activations. However, using task activations from only a single run cut the amount of data contributing to the estimates in half, likely reducing the repeat reliability of the activation estimates. This resulted in lower prediction accuracies than most other analyses (though they remained statistically significant relative to chance).

Each subject’s second task run activations were predicted using activity flow mapping with five distinct sources of FC as input. Note that each subject’s second task run activations were used as input across all FC variants, such that only FC was altered across the five forms of prediction. The number of time points contributing to each FC estimate was matched to the number of time points in each task condition. First, GroupRest involved predicting using a randomly assigned (without replacement) subject’s resting-state FC. It was important to use a single other subject’s FC rather than group-averaged FC here because group-averaged FC would have the unfair advantage (in terms of FC estimation accuracy) of having more data for FC estimation. Further, this approach was more analogous to the Gratton et al. (2018) approach this set of analyses is based on. Second, GroupTask involved using a randomly assigned (without replacement) subject’s task-state FC from the same task as the to-be-predicted task activations. Third, IndivRest used the to-be-predicted subject’s resting-state FC. Fourth, IndivTask used task-state FC estimated from the first task run from the to-be-predicted subject’s data. Thus, task-state FC was estimated from a distinct brain state (run one) from the to-be-predicted task activations (run two). Fifth, IndivTaskRun used task-state FC estimated from the second task run from the to-be-predicted subject’s data. Thus, task-state FC was estimated from the same brain state (run two) as the to-be-predicted task activations (also run two). Predictions of task activations were compared across these five sources of FC to make inferences about the likely contributions of each form of FC to activity flow processes.

### Code availability

All analyses used the publicly available Brain Activity Flow (“Actflow”) Toolbox (https://colelab.github.io/ActflowToolbox/). The code used to call functions in the Actflow Toolbox and run specific analyses will be publicly released upon acceptance of the manuscript for peer-reviewed publication.

## Results

### Task-state FC better models task-evoked activity flow than resting-state FC

We previously found that task-state FC across a variety of tasks differs only minimally from resting-state FC (Cole et al., 2014). Others have replicated this effect (Krienen et al., 2014; Gratton et al., 2018). Further, we recently found that task-state FC correlation strengths are consistently lower than resting-state FC among cortical regions (**Figure 4**), likely driven by task-related local inhibition causing variance and covariance quenching (**Figure 2**) (Ito et al., 2020a). These results seemed to suggest that task-state FC might contribute only minimally to task-related functionality. Consistent with this conclusion, we found that task-evoked activation patterns could be accurately predicted without task-state FC information, using only estimated activity flow over resting-state functional network architecture (Cole et al., 2016; Ito et al., 2017). Here we sought to quantify the contribution of task-state FC to task-evoked processes by directly comparing the prediction of task-evoked activations using activity flow over task-state FC versus activity flow over resting-state FC.

**Figure 4.**
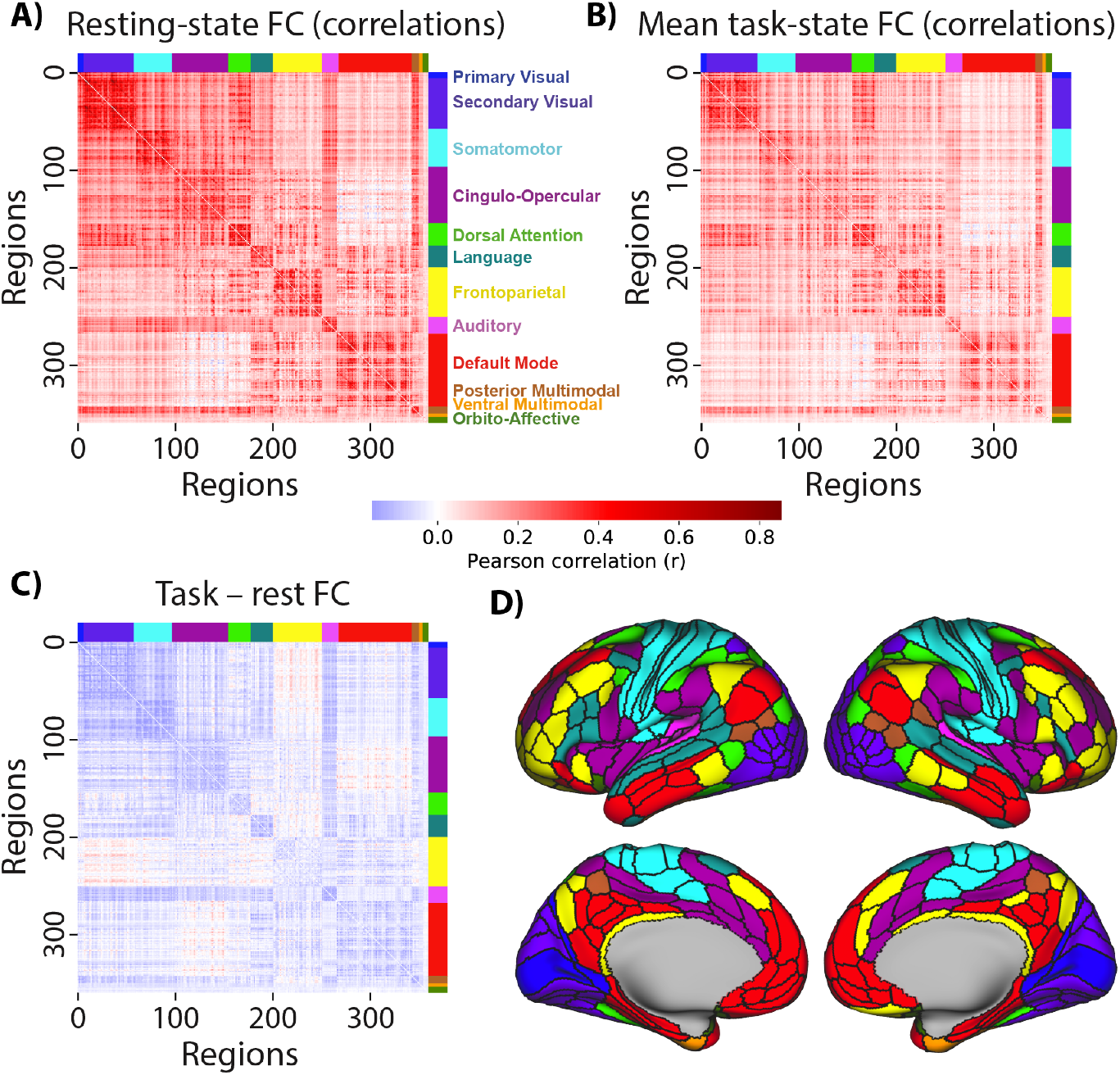
Resting-state and task-state FC are similar and mostly decrease from rest to task. **A**) Resting-state FC correlations averaged across N=176 subjects (discovery set). Network names are listed on the right. **B**) Mean task-state FC correlations averaged across the 24 task conditions. Plotted on the same scale as in A. The similarity of the connectivity matrices in A and B (upper triangle) was r=0.90. **C**) Subtraction between data plotted in panels A and B, plotted on the same scale as in A. Decreased FC was apparent between all networks, with the exception of the fronto-parietal network (which had widespread small increases) and between the cingulo-opercular and default-mode networks. **D**) A cortical surface plot of the networks listed in panel A (Ji et al., 2019) (available at https://github.com/ColeLab/ColeAnticevicNetPartition).

We began by characterizing cortex-wide changes in region-to-region correlations from resting-state FC to task-state FC (**Figure 4C**), focusing for now on cross-task average task-state FC. We found that 68% of connections differed significantly between rest and task (p<0.05, family-wise error corrected for multiple comparisons). Only 4.4% of connections significantly increased from rest to task, while 63.4% of connections significantly decreased from rest to task. Thus, a significantly changed connection was 14 times (63.4 / 4.4 = 14.4) more likely to have decreased than increased from rest to task.

We next carried out an additional replication test, focusing on testing the original activity flow over resting-state FC results (Cole et al., 2016). The original results also used the HCP dataset, but involved fewer subjects (here N=176, before N=100), a less validated preprocessing stream (see Methods), did not include a replication cohort (N=176), and focused on average activation across 7 tasks rather than the 24 conditions included in those tasks. Further, we developed some important innovations relative to that original study: 1) a new set of networks (Ji et al., 2019), 2) utilization of a better-validated set of regions (Glasser et al., 2016), and 3) a focus on predicting a neural population’s “response profile” (condition-wise prediction), rather than focusing solely on whole-brain activation patterns. This final innovation can provide characterization of population receptive fields – a central concept in neuroscience (Wandell and Winawer, 2015) – given that population receptive fields (i.e., the set of stimuli and task conditions that illicit a response in a neural population) can be inferred from condition-wise activity patterns.

As in the previous study, task-evoked activation patterns were predicted (using activity flow estimated over resting-state Pearson correlation FC) with above-chance correspondence between predicted and actual activation patterns: r=0.51, t(175)=99, p<0.00001 (**Figure 5A**). This was true in the replication dataset as well (r=0.51, t(175)=91, p<0.00001), as well as for each of the 24 conditions separately (each p<0.00001). Condition-wise response profiles were also predicted above chance (each p<0.05, FWE corrected) for 100% of the 360 cortical regions analyzed here (**Figure 5D**): mean r=0.52, t(175)=69, p<0.00001 (replication dataset: mean r=0.51, t(175)=71, p<0.00001). These results confirm the informativeness of resting-state FC – which forms the baseline for subsequent tests – for predicting task-evoked activity flow.

**Figure 5.**
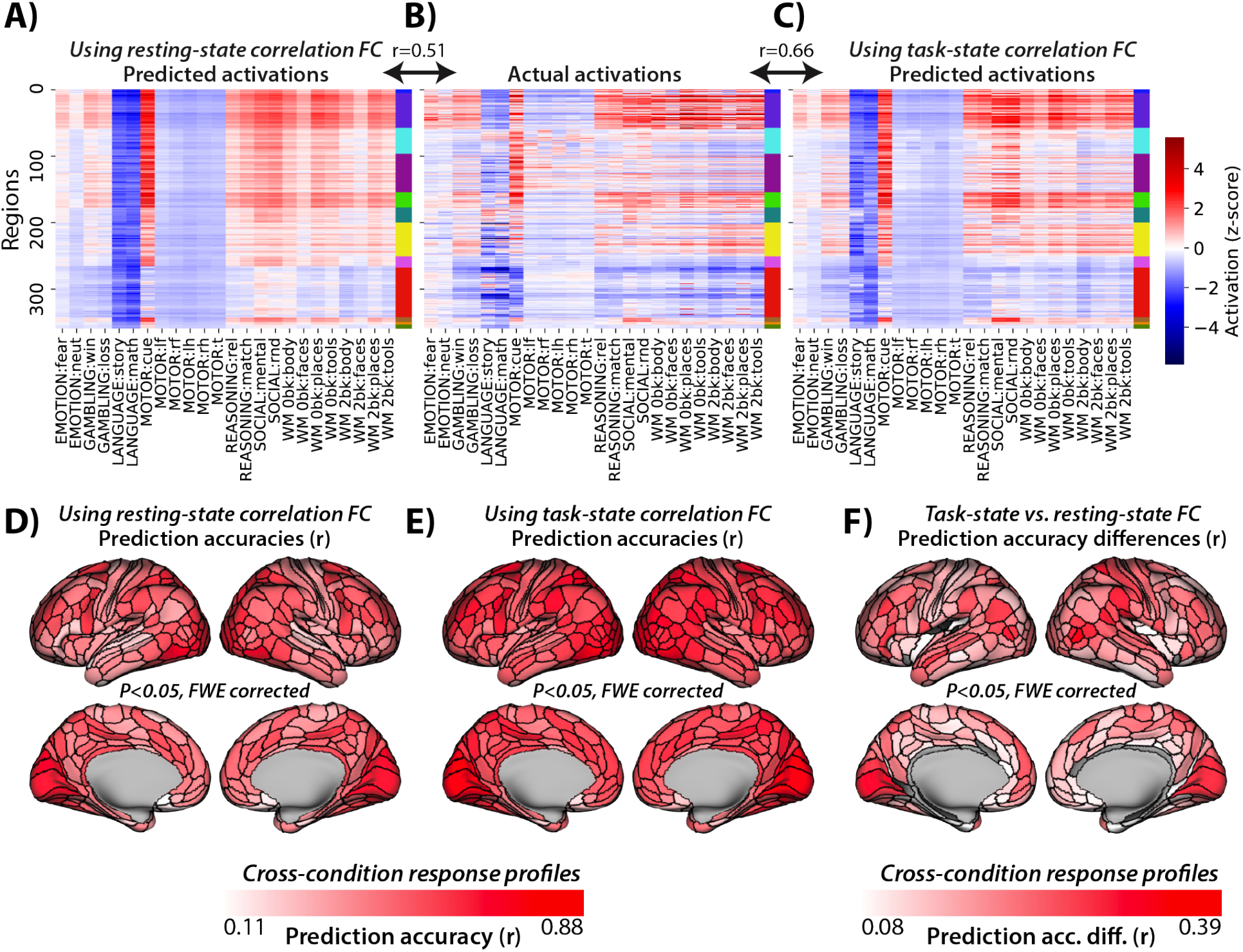
Task-state FC improves correlation-based activity flow models. **A**) Activity flow predictions using resting-state correlation FC across all nodes and conditions (r=0.51 similarity to actual activations, similarity computed for each subject separately prior to averaging r-values). Network colors correspond to those in Figure 4A. **B**) Actual activations (fMRI GLM betas) across all nodes and conditions. **C**) Task-state correlation FC-based activity flow predictions across all nodes and conditions (r=0.66 similarity to actual activations). **D**) Activity flow prediction accuracies using resting-state correlation FC, calculated condition-wise separately for each node. Family-wise error (FWE) corrected for multiple comparisons using permutation testing. All nodes were statistically significant above 0. **E**) Task-state correlation FC-based activity flow prediction accuracies. FWE corrected for multiple comparisons using permutation testing. All nodes were statistically significant above 0. **F**) Task-state vs. resting-state correlation FC-based activity flow prediction accuracy differences. FWE corrected for multiple comparisons using permutation testing. 93% of nodes were statistically significant above 0 (non-significant nodes are gray).

As hypothesized, we found that task-state FC (with task-evoked activations regressed out to avoid circularity in predictions; see Methods) improved activity flow-based predictions of task-evoked activations relative to resting-state FC. Task-state FC was calculated for each task condition, for each subject separately. Activity flow predictions and predicted-to-actual comparisons were also conducted for each task condition and each subject separately (before averaging prediction accuracies across subjects). Activity flow predictions of task activation patterns using task-state correlation FC (**Figure 5C**): r=0.66, t(175)=133, p<0.00001. The direct contrast between activity flow predictions with task-state vs. resting-state FC: r-difference=0.15, t(175)=42, p<0.00001. These results were consistent with the replication dataset (r=0.66, t(175)=123, p<0.00001; r-difference=0.16, t(175)=40, p<0.00001), as well as for each of the 24 conditions separately (each p<0.00001). Condition-wise response profiles were also predicted better (each p<0.05, FWE corrected) with task-state FC than resting-state FC for 93% of the 360 cortical regions analyzed here (**Figure 5F**): mean r-difference=0.12, t(175)=42, p<0.00001 (replication dataset: mean r-difference=0.13, t(175)=38, p<0.00001). These results confirm our primary hypothesis: That task-state FC is more informative regarding the paths of task-evoked activity flow than resting-state FC.

### Prediction accuracy improves when causal confounding is reduced

Multiple regression is a standard statistical method that conditions on other variables, such that multiple regression parameter estimates indicate unique variance contributing to each prediction. Thus, when used for FC estimation with each node (one-at-a-time) being predicted by all other nodes’ time series, multiple regression reduces causal confounds (e.g., a third variable causing a spurious association) in inferred associations (Cole et al., 2016; Ito et al., 2017; Sanchez-Romero and Cole, 2020). We used it here given the goal of improving FC-based causal inferences (though limitations remain even when using multiple-regression FC rather than Pearson-correlation FC) (Reid et al., 2019; Sanchez-Romero and Cole, 2020). We hypothesized that multiple-regression FC would provide more accurate task-evoked activation predictions with activity flow mapping (using both resting-state FC and task-state FC) than when using Pearson-correlation FC, due to reduction in causal confounds.

Prior to testing this hypothesis we analyzed the change in multiple-regression FC values across rest and task (**Figure 6**). We controlled for the amount of data across methods by calculating resting-state FC using the same number of time points as when estimating task-state FC for each of the 24 task conditions. The resulting connectivity coefficients were much smaller and the connectivity matrix was sparser when using multiple-regression FC, consistent with multiple-regression FC reducing the number of confounds relative to correlation-based FC (Reid et al., 2019; Sanchez-Romero and Cole, 2020). We then averaged across these 24 FC estimates for analysis and visualization. The mean resting-state FC and task-state FC were highly similar to each other (r=0.94), as is the case for correlation-based FC (Cole et al., 2014). We found that 2.6% of connections (915 connections) differed significantly between rest and task (p<0.05, family-wise error corrected for multiple comparisons). Only 0.28% of connections (180 connections) significantly increased from rest to task, while 2.28% of connections (1478 connections) significantly decreased from rest to task. Thus, a significantly changed connection was eight times as likely to have decreased than increased from rest to task. The small FC differences for task-state FC relative to resting-state FC (even smaller than with Pearson-correlation FC) might make it surprising if using multiple-regression task-state FC improves activity flow prediction accuracy. Yet we hypothesized that the increase in activity flow prediction accuracy would remain and perhaps even be enhanced given that activity flows sum across many sources, potentially allowing for large differences in activation prediction values despite small differences in contributing connectivity values.

**Figure 6.**
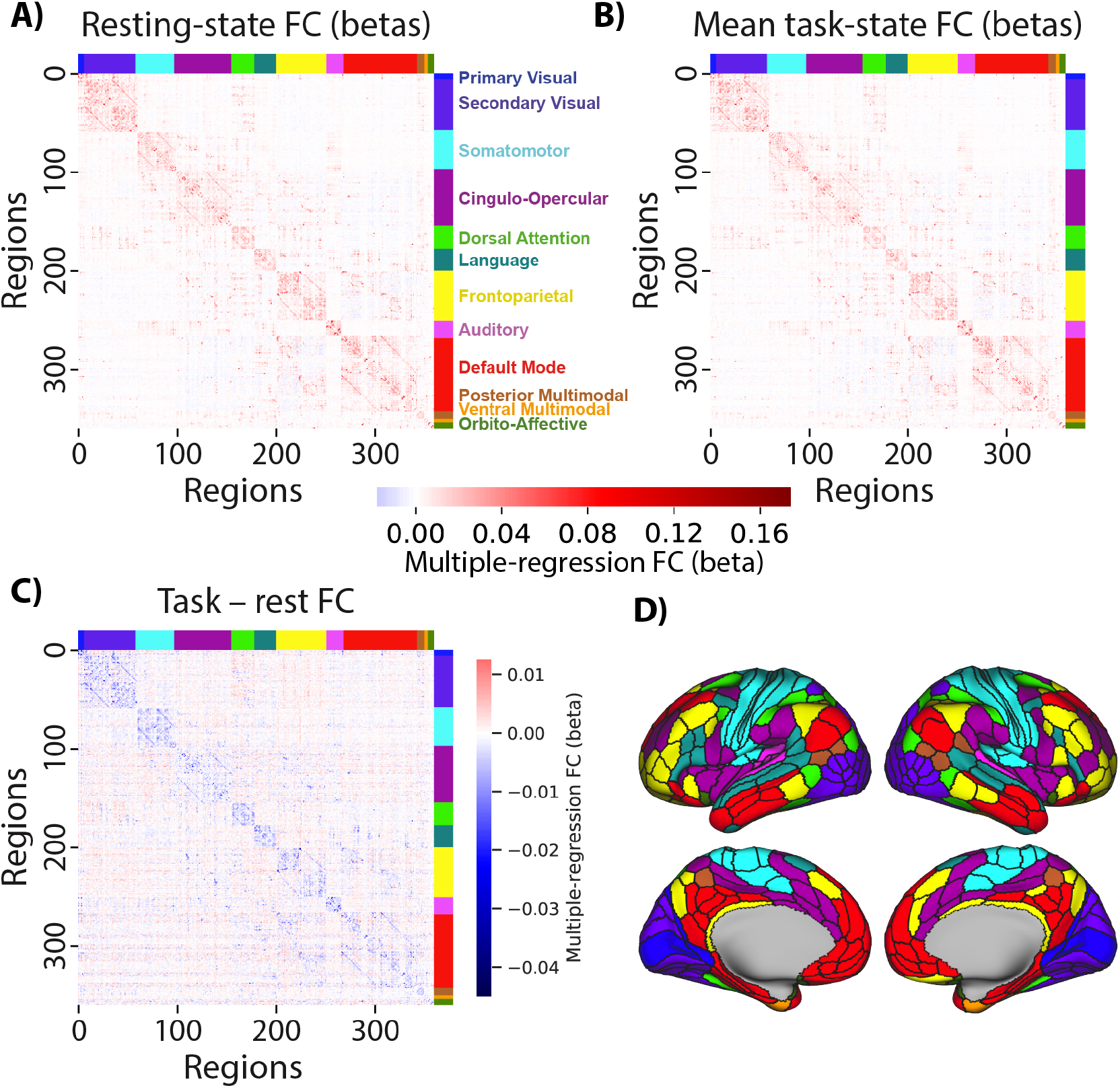
Multiple-regression FC is similar across rest and task and mostly decreases from rest to task. **A**) Resting-state multiple-regression FC averaged across N=176 subjects (discovery set). Network names are listed on the right. **B**) Mean task-state multiple-regression FC averaged across the 24 task conditions. The similarity between the matrices shown in panels A and B was r=0.94. **C**) Subtraction between data plotted in panels A and B. Only 2.6% of connections were statistically different from 0. **D**) A cortical surface plot of the networks listed in panel A (Ji et al., 2019) (available at https://github.com/ColeLab/ColeAnticevicNetPartition).

Using multiple-regression FC with resting-state data, task-evoked activation patterns were again predicted with above-chance correspondence between predicted and actual activation patterns: r=0.46, t(175)=81, p<0.00001 (**Figure 7A**). This was true for each of the 24 conditions separately (each p<0.00001). Condition-wise response profiles were also predicted above chance (each p<0.05, FWE corrected) for 98% of the 360 cortical regions analyzed here (**Figure 7D**): mean r=0.46, t(175)=99, p<0.00001. Note that the number of time points used for the task-state FC results were matched for the estimation of resting-state FC (see Methods).

**Figure 7.**
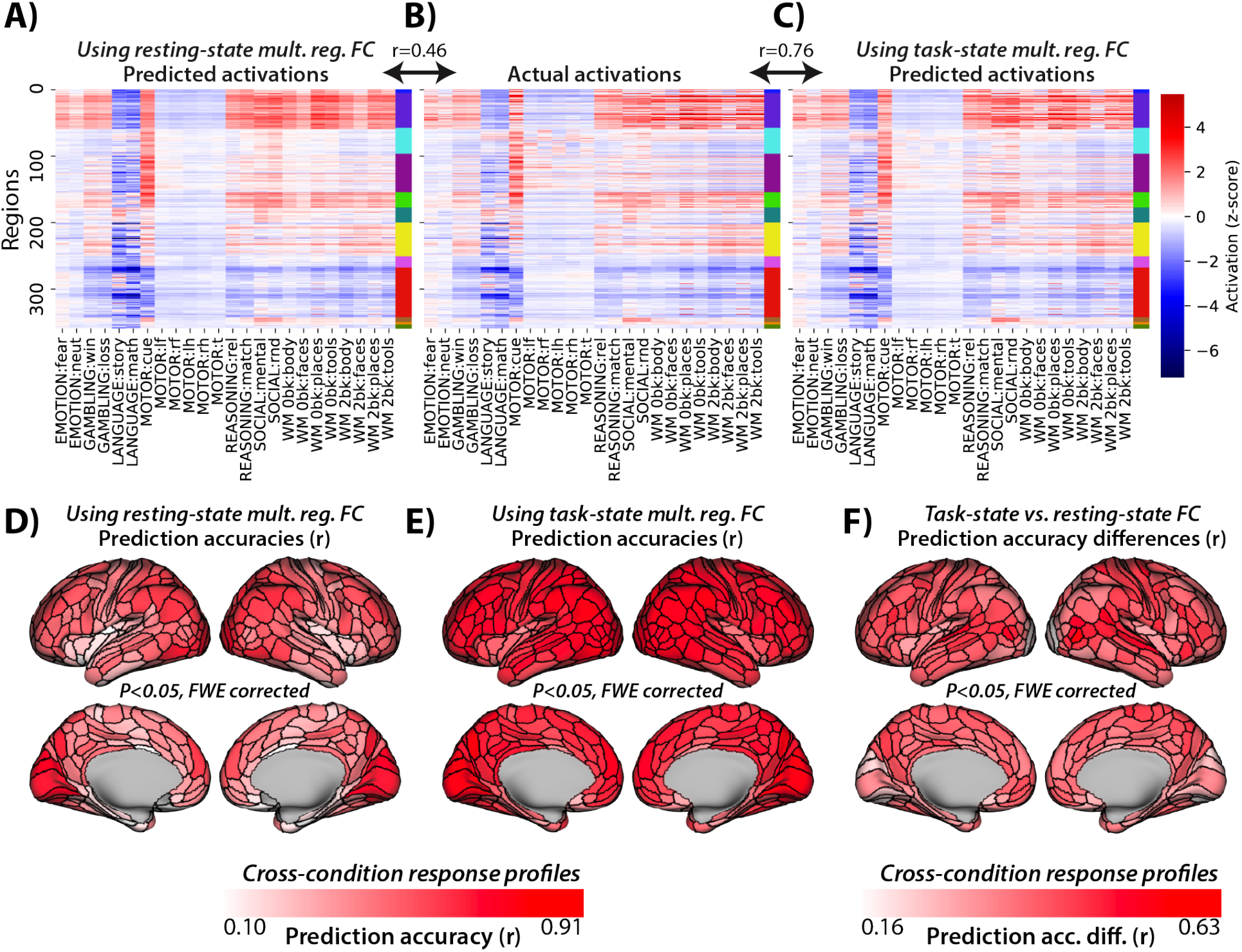
Task-state FC improves multiple-regression-based activity flow models. **A**) Resting-state multiple-regression FC-based activity flow predictions across all nodes and conditions (r=0.46 similarity to actual activations, similarity computed for each subject separately prior to averaging r-values). Network colors correspond to those in Figure 6A. **B**) Actual activations (fMRI GLM betas) across all nodes and conditions. **C**) Task-state multiple-regression FC-based activity flow predictions across all nodes and conditions (r=0.76 similarity to actual activations). **D**) Resting-state multiple-regression FC-based activity flow prediction accuracies, calculated condition-wise separately for each node. FWE corrected for multiple comparisons using permutation testing. 98% of nodes were statistically significant above 0. **E**) Task-state multiple-regression FC-based activity flow prediction accuracies. FWE corrected for multiple comparisons using permutation testing. All nodes were statistically significant above 0. **F**) Task-state vs. resting-state multiple-regression FC-based activity flow prediction accuracy differences. FWE corrected for multiple comparisons using permutation testing. All nodes were statistically significant above 0.

As hypothesized, we also found with multiple-regression FC that task-state FC improved activity flow-based predictions of task-evoked activations relative to resting-state FC. Activity flow predictions of task activation patterns using task-state multiple-regression FC (**Figure 7C**): r=0.76, t(175)=146, p<0.00001. The direct contrast between activity flow predictions with task-state vs. resting-state FC: r-difference=0.31, t(175)=94, p<0.00001. These results were consistent with the replication dataset (r=0.77, t(175)=149, p<0.00001; r-difference=0.31, t(175)=95, p<0.00001), as well as for each of the 24 conditions separately (each p<0.00001). Condition-wise response profiles were also predicted better (each p<0.05, FWE corrected) with task-state FC better than resting-state FC for 100% of the 360 cortical regions analyzed here (**Figure 7F**): mean r=0.75, t(175)=148, p<0.00001. The direct contrast between task-state FC and resting-state FC-based condition-wise predictions: mean r-difference=0.27, t(175)=106, p<0.00001. These results further confirm our primary hypothesis: That task-state FC is more informative regarding the paths of task-evoked activity flow than resting-state FC. A summary of these and related results is included in **Table 2**.

**Table 2.**
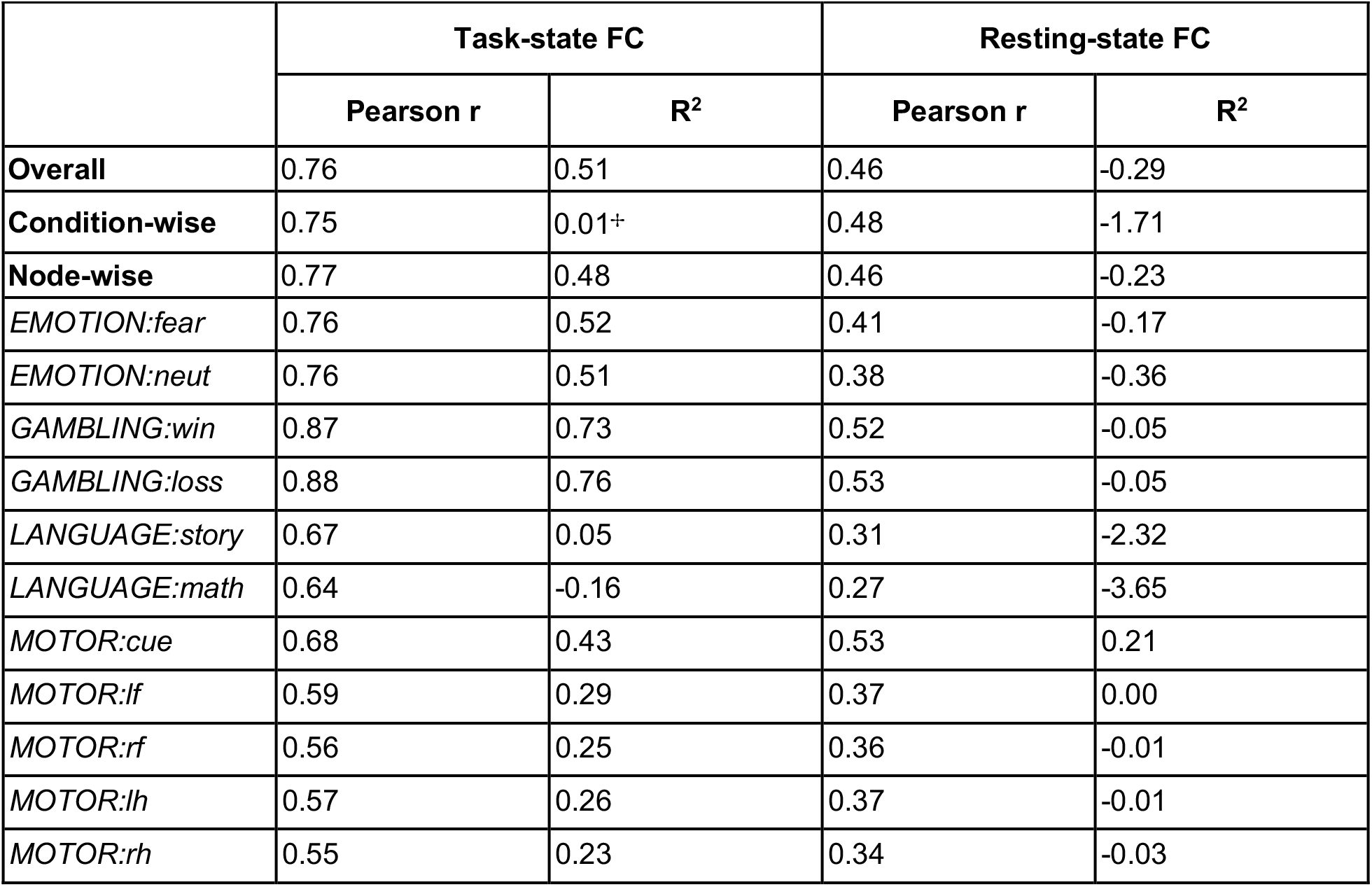
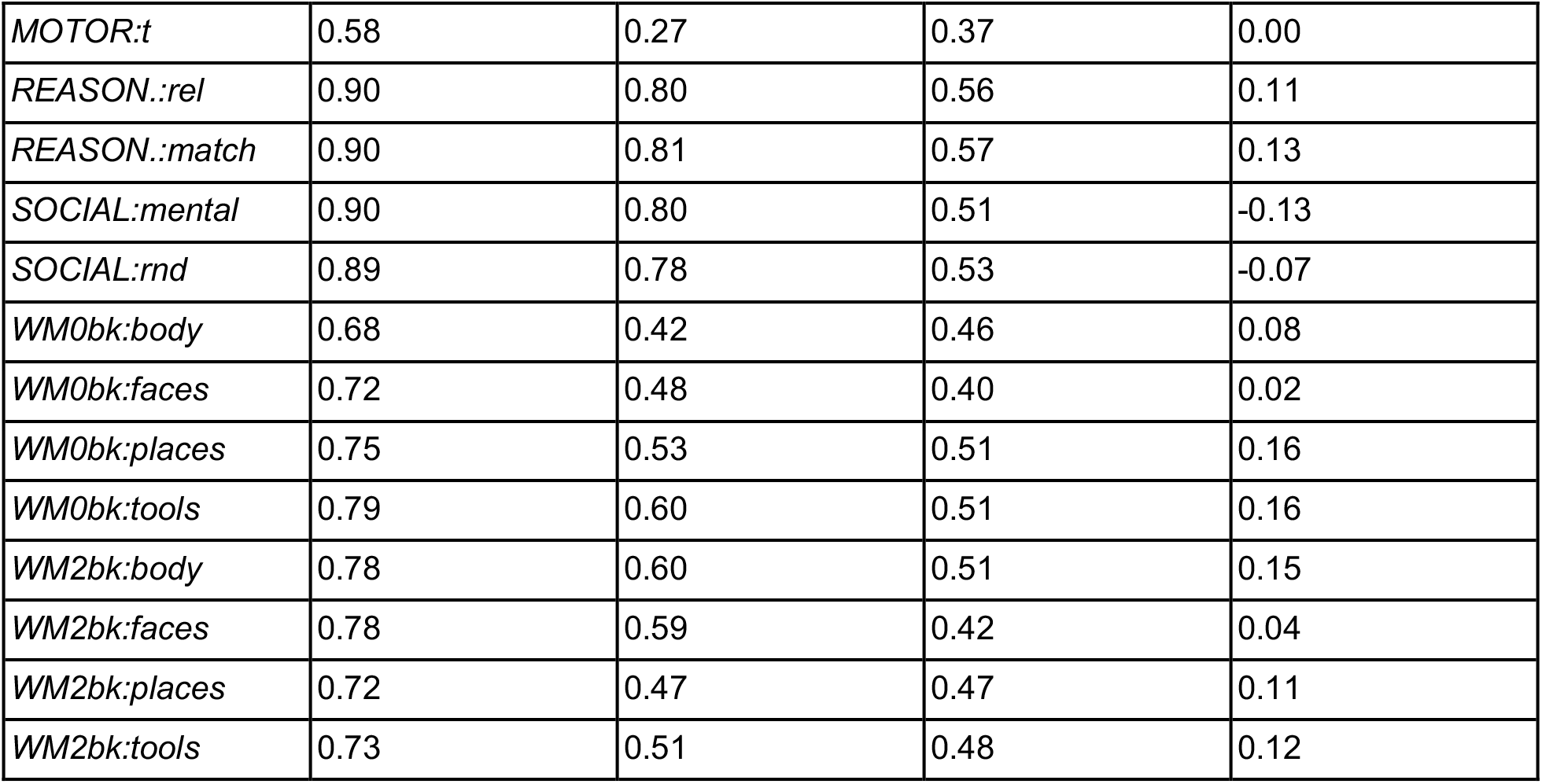
Summary of Pearson r and R^2^ results. Pearson r and R^2^ prediction accuracy assessment results are reported across task-state FC and resting-state FC. Results comparing all predicted to actual activation values at once are included (“overall”), as well as results comparing prediction accuracy across conditions for each node separately (“condition-wise”) and prediction accuracy across nodes for each condition separately (“node-wise”). See Table 1 for descriptions of Pearson r vs. R^2^ and condition-wise vs. node-wise predictions. Node-wise comparisons for each task condition separately are included in the last 24 rows. ^✢^Mean condition-wise task-state FC R^2^ values were dragged down by regions with especially poor predictions due to misspecified scales. Since misspecified scales affect R^2^ but not Pearson r-values, poor scaling of predictions can be detected when R^2^ predictions are much worse than Pearson r predictions. We found that 38 brain regions had R^2^ values below −1.0, meaning the predictions were 100% worse than the average activation across all task conditions for those regions. Notably, these same regions all had above-0 Pearson r-values (mean r=0.43), demonstrating this was primarily a scaling issue. In contrast, 186 (of 360) brain regions had R^2^ above 0.25, such that most brain regions exhibited overall accurate condition-wise predictions even when taking scale into account.

|  | Task-state FC |  | Resting-state FC |  |
| --- | --- | --- | --- | --- |
| | Pearson $r$ | $R^2$ | Pearson $r$ | $R^2$ |
| **Overall** | 0.76 | 0.51 | 0.46 | -0.29 |
| **Condition-wise** | 0.75 | 0.01+ | 0.48 | -1.71 |
| **Node-wise** | 0.77 | 0.48 | 0.46 | -0.23 |
| *EMOTION:fear* | 0.76 | 0.52 | 0.41 | -0.17 |
| *EMOTION:neut* | 0.76 | 0.51 | 0.38 | -0.36 |
| *GAMBLING:win* | 0.87 | 0.73 | 0.52 | -0.05 |
| *GAMBLING:loss* | 0.88 | 0.76 | 0.53 | -0.05 |
| *LANGUAGE:story* | 0.67 | 0.05 | 0.31 | -2.32 |
| *LANGUAGE:math* | 0.64 | -0.16 | 0.27 | -3.65 |
| *MOTOR:cue* | 0.68 | 0.43 | 0.53 | 0.21 |
| *MOTOR:lf* | 0.59 | 0.29 | 0.37 | 0.00 |
| *MOTOR:rf* | 0.56 | 0.25 | 0.36 | -0.01 |
| *MOTOR:lh* | 0.57 | 0.26 | 0.37 | -0.01 |
| *MOTOR:rh* | 0.55 | 0.23 | 0.34 | -0.03 |
| *MOTOR:t* | 0.58 | 0.27 | 0.37 | 0.00 |
| *REASON.:rel* | 0.90 | 0.80 | 0.56 | 0.11 |
| *REASON.:match* | 0.90 | 0.81 | 0.57 | 0.13 |
| *SOCIAL:mental* | 0.90 | 0.80 | 0.51 | -0.13 |
| *SOCIAL:rnd* | 0.89 | 0.78 | 0.53 | -0.07 |
| *WM0bk:body* | 0.68 | 0.42 | 0.46 | 0.08 |
| *WM0bk:faces* | 0.72 | 0.48 | 0.40 | 0.02 |
| *WM0bk:places* | 0.75 | 0.53 | 0.51 | 0.16 |
| *WM0bk:tools* | 0.79 | 0.60 | 0.51 | 0.16 |
| *WM2bk:body* | 0.78 | 0.60 | 0.51 | 0.15 |
| *WM2bk:faces* | 0.78 | 0.59 | 0.42 | 0.04 |
| *WM2bk:places* | 0.72 | 0.47 | 0.47 | 0.11 |
| *WM2bk:tools* | 0.73 | 0.51 | 0.48 | 0.12 |

These multiple-regression FC prediction accuracies were higher than the correlation FC prediction accuracies. We quantified this improvement by comparing these predictions accuracies statistically. The improvement in overall prediction accuracy across all nodes and conditions using task-state FC with multiple-regression FC (relative to correlation FC) was highly significant: mean r-difference=0.10, t(175)=34, p<0.00001. This suggests that the influence of task-state FC changes on task-evoked activations is more accurately described when accounting for causal confounds using multiple-regression FC (Reid et al., 2019; Sanchez-Romero and Cole, 2020).

### Visualizing predictions across diverse cognitive domains on brain surfaces

Three of the 24 task conditions were selected for detailed illustration due to the diversity of cognitive demands they represented (**Figure 8**). As with all other task conditions, these three task conditions were significantly better predicted using task-state FC than resting-state FC (all p<0.00001). To reduce redundancy with Figure 7, we used a different metric to quantify prediction accuracy: average-then-compare R^2^ prediction accuracies. This involves averaging activations across subjects prior to comparing predicted and actual activation patterns, then quantifying similarity using R^2^ scores (see **Table 1** for details). R^2^ scores were used such that the scale of the values was taken into account, not just the unscaled pattern similarity quantified using Pearson correlations. Average-then-compare was used so predicted-to-actual similarity reflected the similarity of the group-averaged values visualized in Figure 8. However, note that it is more accurate to compare at the individual-subject level prior to group averaging the similarity values, as was done for all other analyses. This is due to individual-specific activations and FC being taken into account, rather than blurring activity and connectivity patterns across individuals prior to predicted-to-actual comparison.

**Figure 8.**
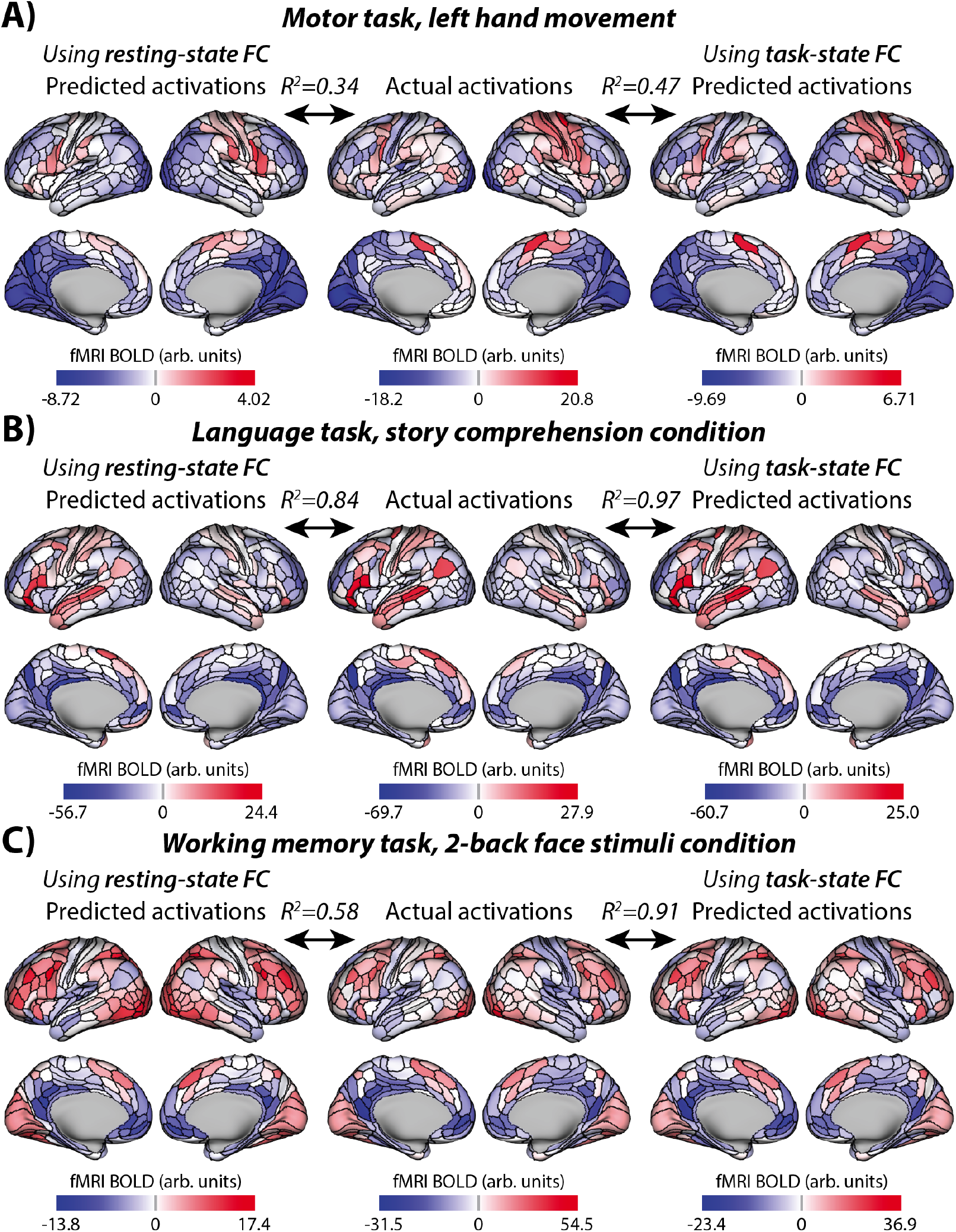
Visualizing three example task conditions across diverse cognitive domains. These values are all present in Figure 7 – three of the columns from the matrices shown in Figure 7A, 7B, and 7C – but are visualized here on cortical anatomy. R^2^ values are based on the average-then-compare approach, quantifying the similarity of what is being visualized (i.e., group-level rather than individual-level similarity). The average-then-compare approach resulted in higher accuracies than the compare-then-average approach used in Table 2 (and elsewhere); see Table 1. **A**) Activations for the motor task, left hand movement condition is shown. Note the right somatomotor activations corresponding to the left hand movement, which is most prominent for the actual and task-state FC-based predictions. Also note the scale difference for resting-state FC predictions, which contributes to R^2^ (not Pearson correlation) values. **B**) Activations for the language task, story comprehension condition. Note the left-lateralized language network activation pattern in all three maps. **C**) Activations for the working memory task, 2-back face stimuli condition. Note the larger improvement in group-level prediction accuracy in this task condition relative to the motor and language conditions. It will be important for future research to identify factors underlying these task condition differences.

### The role of task-state FC decreases versus increases in activity flow predictions

A potentially counter-intuitive aspect of task-state FC estimates is their tendency to decrease relative to resting-state FC estimates (see **Figure 2**). Here we sought to determine whether these decreases are functionally meaningful. Given that most statistically significant task vs. rest FC changes decreased from rest to task, and yet task-state FC improved activity flow prediction accuracy overall, it is likely that the decreases played a prominent role in improving activity flow prediction accuracies. We sought to better establish this possibility using a lesioning approach – selectively removing connections of interest so as to quantify their importance for prediction accuracy.

Task-state multiple-regression functional connections that decreased from rest were lesioned (on a subject-specific and task-specific basis) by setting those connections to 0. We then conducted activity flow mapping with all 360 nodes and 24 task conditions. Using FC-decrease lesioned models we found that prediction accuracy reduced to r=0.48 (from r=0.76 without lesioning): r-difference=0.29, t(175)=59, p<0.00001. This suggests that rest-to-task FC decreases were important for task activation prediction, consistent with the larger number of rest-to-task FC decreases relative to increases.

We also found that task-state FC increases from rest were functionally important. Using FC-increase lesioned models we found that prediction accuracy reduced to r=-0.20 (from r=0.76 without lesioning): r-difference=0.96, t(175)=87, p<0.00001. Further, the FC-increase lesioned model performed significantly worse than the FC-decrease mode (r=0.48 vs. r=-0.20): r-difference=0.67, t(175)=50, p<0.00001. This result confirmed the functional importance of rest-to-task increases in FC, while also revealing that FC increases have a larger impact of task activation prediction accuracy than FC decreases. This is despite the smaller number and amplitude of statistically significant increases relative to decreases.

Note, however, that we did not restrict our lesioning to statistically significant FC changes. Rather, all connections were considered so as to avoid biasing results to only those FC changes that are consistent across subjects. An analysis of the connections lesioned in these two models revealed that 50.2% of connections decreased from rest to task, while 49.8% increased. Further, FC decreases changed by 0.461 on average, while FC increases changed by 0.462 on average. These results revealed that – once results were not restricted to statistically significant group-level FC changes – FC increases were nearly as numerous and large in amplitude as FC decreases.

The large number of task-state FC decreases (just over half of FC changes) may appear counterintuitive due to the common intuition that as activity increases during task performance so too should FC, reflecting an increase in neural interactions during tasks. We hypothesized that this intuition might more accurately apply to activity flows (*activations* ✕ *FC*), rather than FC alone. Specifically, we expected that a large portion of activity flows would be positive – possibly reflecting increases in neural interactions – despite restricting to only FC decreases from resting state. This is possible because activity flows reflect activations (changes in activity levels from inter-block rest baseline) multiplied by connectivity, such that even FC decreases (e.g., 0.25, down from 0.40) can be multiplied by a positive activation increase from resting baseline (e.g., 0.30) to yield a positive activity flow (e.g., 0.25 ✕ 0.30 = 0.075) (see **Figures 1C & 1D**). As expected, we found that 50.001% of all non-zero activity flows (over task-reduced connections) were positive. These results suggest that a task-state FC decrease can nonetheless result in increased activity in a distal region (via activity flow), despite activity increases in one region having a smaller effect on the other (relative to resting state).

### The role of individual-specific factors in task-state FC

To this point the task-state FC advantage has been shown when activity flow predictions are computed in a subject-specific manner. Further, task-state FC has been computed using the same task runs as the to-be-predicted task activations. We next sought to quantify the impact of subject-specific (vs. group) FC and run-specific (vs. FC from an independent run) FC. Confirming the importance of these factors, a recent study of group vs. individual-subject FC indicated there are large effects of subject-specific FC, and also of subject-specific task-state FC relative to subject-specific resting-state FC (Gratton et al., 2018). That study quantified these effects by comparing whole-cortex connectivity patterns across individuals and cognitive states, such as assessing how similar the same subject’s vs. other subjects’ task-state FC patterns were. They found high similarity between resting-state FC and task-state FC connectivity matrices within subject (relative to between subjects), indicating a large effect of subject-specific resting-state FC on specifying each subject’s task-state FC connectivity pattern. They also found that these subject-specific effects were less prominent (relative to group effects) for task activations (i.e., comparing whole-cortex activation patterns) relative to FC. Since activity flow mapping is a combination of both task activations and FC the relative role of subject-specific vs. group effects in activity flow predictions is unclear.

Based on Gratton et al. (2018), group FC was defined as other subjects’ FC. Thus, the effect of group (non-individualized) FC was quantified by using a randomly assigned (without replacement) subjects’ FC for each subject’s activity flow prediction. It was important to use other individual subjects’ FC – rather than group-averaged FC – because it was important to control for the total amount of data going into the FC estimates when comparing group vs. individualized FC effects.

We compared the effects of three factors: group vs. individual, rest vs. task, and same vs. different task run. This revealed statistically significant (all p<0.00001) effects for all three factors, showing increases for individual, task, and same-run factors (**Figure 9**). This was the case for both Pearson correlation (**Figure 9A**) and R^2^ (**Figure 9B**) assessments of activity flow prediction accuracy. The largest effect on activity flow prediction accuracies was from individualized task-state FC (**Figure 9C**). This effect (R^2^=0.40) was several times larger than the effect of individualized rest/intrinsic FC (R^2^=0.13), which can be thought of as a baseline (e.g., due to lower overall metabolic demand in the brain during resting state (Gusnard et al., 2001)). This suggests (consistent with Gratton et al. (2018)) that individualized task-state FC reconfigurations play a large role in shaping task activations. Note that the individualized nature of task-state FC may help explain why there is a large effect of task-state FC on activity flow prediction accuracy despite there being only small group-level differences in task-state FC relative to resting-state FC (r=0.94 similarity between group-level resting-state and task-state FC matrices; see **Figure 6A & 6B**).

**Figure 9.**
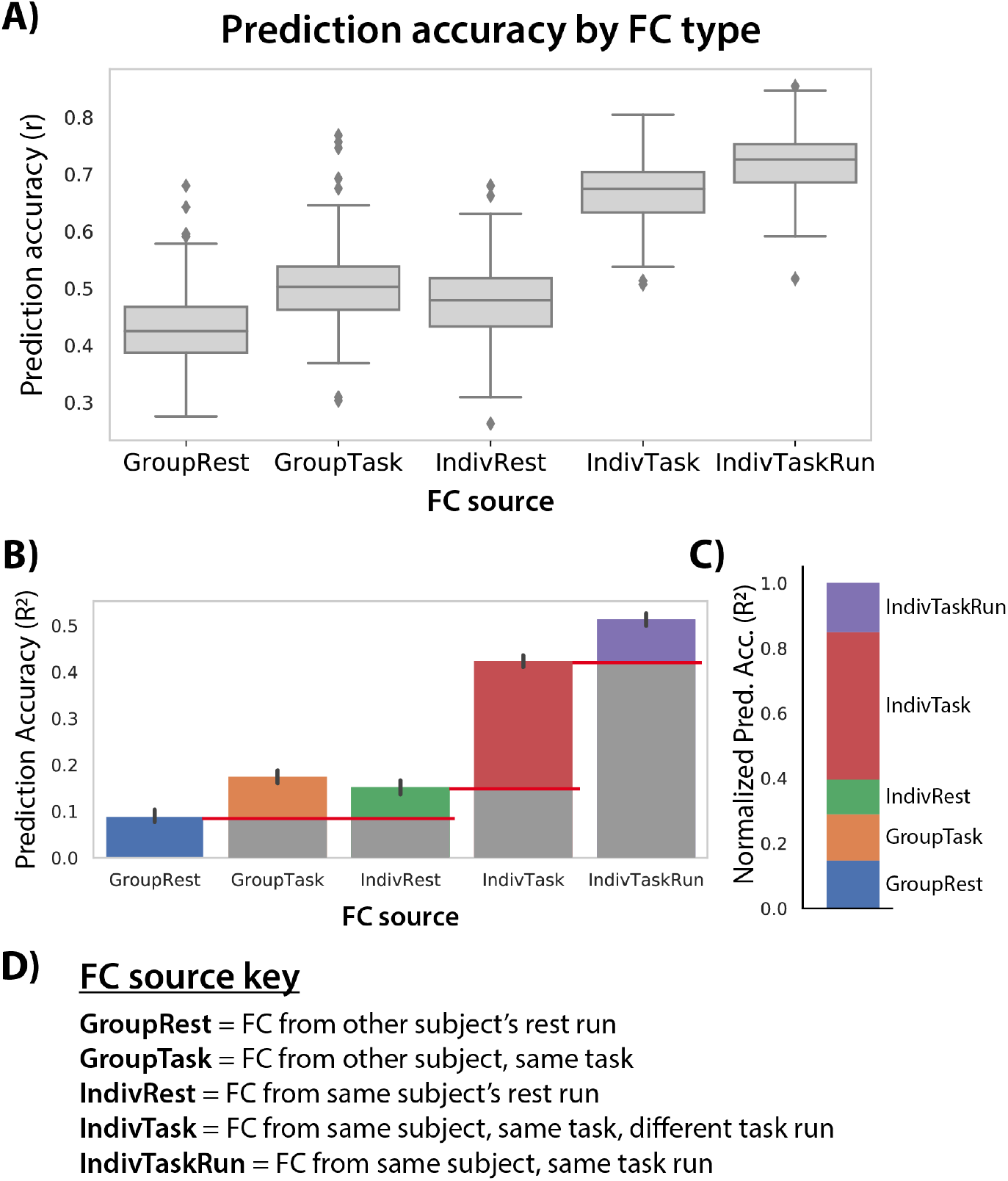
Intrinsic, individual, task-specific, and task-run-specific factors contribute to shaping each individual’s task activations. **A**) Distinct sources of FC were used for activity flow mapping with the same task activations (24 conditions, first run only). The prediction accuracies can be directly compared given that they are predicting the same set of activations (prediction accuracy computed across 360 regions and 24 conditions). **B**) Prediction accuracies computed as unscaled R^2^ (not r^2^), which can be interpreted (once multiplied by 100) as percent of variance in the to-be-predicted activity pattern explained by the prediction. The red lines indicate baselining to quantify relative effects in panel C. **C**) Same as panel B, but baselined (as indicated in panel B) to highlight relative effects and normalized such that all values add up to 1. This facilitates interpretation of the role of intrinsic, individual, task, and task-run-specific factors in producing task activations. These normalized R^2^ values can be interpreted as the proportion of the explained variance (in the IndivTaskRun results) contributed to by each factor. **D**) Detailed descriptions of the FC sources.

A notable discrepancy in the results is the consistently higher R^2^ values in Figure 9B compared to Table 2, especially for the resting-state-FC-based predictions (−0.29 in Table 2 versus over 0.1 in Figure 9B). This is despite half the data used per prediction in Figure 9B (one run per task) relative to Table 2 (two runs per task). We hypothesized this was due to overfitting to noise in the multiple-regression FC estimates, which was likely substantially reduced in the Figure 9B analysis due to the use of fewer principal components (greater regularization) in the principal components regression step. This reduction in the included number of principal components was by necessity (given fewer data points than regressors; see Methods), but may have nonetheless improved the generalization of the regression weights by removing noise variance from the data prior to the multiple regression fitting step. Confirming this, we found that R^2^ increased from −0.29 to 0.38 when everything was identical to the analysis in Table 2 (resting-state FC overall R^2^) except that half of the principal components were used (matching the number of principal components used in Figure 9B). This demonstrates that exploring the role of regularization in activity flow-based predictions will be important for future research (e.g., using cross-validation to optimize the number of principal components). Importantly, using this more optimal regularization also improved the task-state-FC-based prediction from 0.51 (in Table 2) to 0.60. This demonstrates that regularization also improves task-FC-based predictions. Note that the low average R^2^ performance for condition-wise task-state-FC-based prediction (see Table 2) was also improved with the increased regularization (from R^2^=0.01 to R^2^=0.19).

### Task-general FC also improves prediction accuracy

We next sought to determine whether task-state FC estimates needed to come from the same task as the to-be-predicted task activations in order to improve predictions. Instead, it might be that any (or most) task states provide a similar advantage relative to resting-state FC. This would be consistent with observations that many task-state FC changes from resting-state FC generalize across task states (Cole et al., 2014; Schultz and Cole, 2016). Further, using task-general FC (connectivity estimated by concatenating time series from multiple task states) could have a benefit to prediction accuracy based on the increased number of time points going into the FC estimates, given limited data from any single task. We computed task-general FC simply as multiple regression coefficients calculated across all task runs. We avoided circularity by removing mean evoked responses from each task condition (as with task-specific FC estimation), as well as by excluding the time points during the to-be-predicted task condition. Thus, the inference was similar to using resting-state FC for activity flow mapping, since the connectivity models were based on a brain state independent of the to-be-predicted task condition.

Consistent with our hypothesis that task-general FC carries some of the same benefits as task-state FC, we found that using task-general FC increased prediction accuracy relative to using resting-state FC (matched for the number of time points). The overall prediction accuracy (across all nodes and task conditions) with task-general FC was r=0.90 (t(175)=156, p<0.00001), while the same analysis with resting-state FC yielded r=0.87 (t(175)=145, p<0.00001). Comparing the two approaches, the mean r-difference=0.03 (t(175)=33, p<0.00001). Condition-wise response profiles were also predicted significantly above chance with task-general FC (mean r=0.89, t(175)=167, p<0.00001) and resting-state FC (mean r=0.87, t(175)=27, p<0.00001). These condition-wise response profile predictions were significantly better (each p<0.05, FWE corrected) with task-general FC than resting-state FC for 41% of the 360 cortical regions: mean r-difference=0.02, t(175)=27, p<0.00001. These results demonstrate a small but reliable benefit to using task (rather than rest) data to estimate FC, even if the task data come from task conditions other than the one being modeled.

It is possible that not all task conditions improved activity flow prediction accuracy for every other task condition. For instance, two very distinct tasks might actually bias task-state FC away from the correct relationship between brain regions during each task (relative to resting-state FC). We tested this possibility by iteratively using each task condition’s task-state FC for predicting whole-cortex brain activity patterns for each task condition. This produced a state-generalization matrix using task-state FC (**Figure 10A**). All task conditions could be used individually to predict all other task conditions (mean r=0.49, t(175)=19, p<0.00001). The same was found using time-matched resting-state FC (mean r=0.45, t(175)=30, p<0.00001) (**Figure 10B**). Overall (averaged across all values in the state-generalization matrices), task-state FC predicted task activations better than resting-state FC: r-difference=0.04, t(176)=4, p=0.0004). However, when each condition pair was compared separately (**Figure 10C**), task-state FC did not better predict all conditions individually. Indeed, there were cases of both significantly (p<0.05, Bonferroni corrected for multiple comparisons) increased and decreased prediction accuracy (**Figure 10D**).

**Figure 10.**
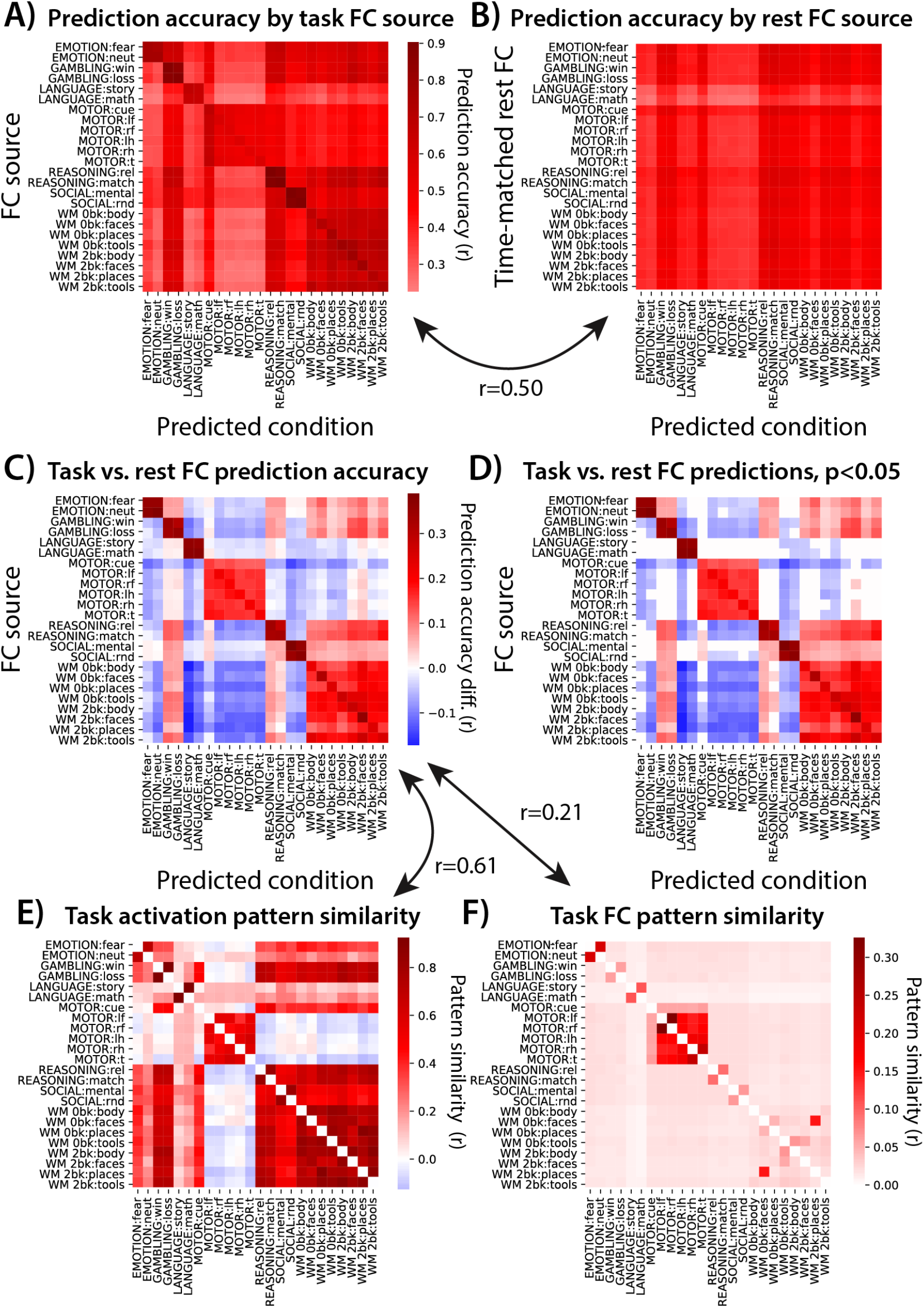
Activity flow routes are better described by similar task conditions. **A**) Generalization of each condition’s task-state FC was tested across each condition’s whole-cortex task activation pattern, quantified by Pearson correlation r-values. All cells of the matrix were significantly higher than 0, p<0.05 Bonferroni corrected for multiple comparisons. **B**) Identical to panel A, but with resting-state FC used instead of task-state FC. The number of time points going into each FC estimate was matched to the task-state FC estimates. Again, all cells of the matrix were significant, p<0.05 Bonferroni corrected. Variation along the y-axis reflects the effect of the amount of time contributing to each FC estimate, while variation along the x-axis reflects predictability of task activation patterns. **C**) Subtraction of the matrix in panel B from the matrix in panel A. **D**) Thresholded version of the matrix in C, p<0.05 Bonferroni corrected. **E**) Similarity of task activation patterns across each pair of task conditions. Pearson correlations using whole-cortex activation patterns. **F**) Similarity of task-state FC patterns across each pair of task conditions. Pearson correlations across whole-cortex multiple-regression FC matrices.

The matched-condition cases (diagonal in the **Figure 10D** matrix) and matched-task cases (conditions from the same task) were universally higher for task-state FC. We further found that the overall pattern of prediction accuracy differences was correlated with whole-cortex task activation pattern similarities (**Figure 10E**; r=0.61, t(175)=95, p<0.00001). This was also the case (but less so) for task-state FC pattern similarities (**Figure 10F**; r=0.21, t(175)=39, p<0.00001). Similarity between tasks was quantified using Pearson correlations, between 360-length vectors (activations for all regions) in Figure 10E and 129,240-length vectors (multiple-regression FC among all regions) in Figure 10F. These results suggest that similar task conditions (quantified as task activation pattern similarity) better describe activity flow routes for each other than dissimilar task conditions. This was further confirmed by comparing predictions between task conditions with positive task activation correlations (mean r=0.041) versus tasks conditions with negative task activation correlations (mean r=-0.046) directly: r-difference=0.087, t(175)=32, p<0.00001.

### The role of amount of data used for FC estimation

For most analyses we restricted the amount of data used for estimating resting-state FC to the amount used for each task-state FC estimate. This controlled for the amount of data as a factor, making for a more fair comparison between the methods. However, there is often more resting-state data available in fMRI datasets than the 114 time points (82 seconds) available on average for each of the 24 task conditions used here. We therefore next used the entire amount of resting-state fMRI data available for estimating resting-state FC: 4780 time points (57 minutes).

As expected, task-evoked activation patterns were better predicted with more resting-state fMRI data used for estimating multiple-regression FC. Predicted-to-actual activation pattern similarity across all nodes and task conditions: r=0.89, t(175)=166, p<0.00001. This was significantly higher than when the amount of data was matched to the task conditions (time-matched): r-difference=0.31, t(175)=94, p<0.00001. The mean R^2^ was 0.78, meaning 78% of the activation pattern variance was accurately predicted on average. This was substantially larger than the R^2^ of −0.29 with the time-matched results. A positive r-value along with a negative R^2^ suggests there was a scaling issue with the time-matched resting-state FC results (see Table 1).

These results demonstrate that much of the predictive advantage of task-state FC is present with low amounts of data. This likely reflects the fact that prediction has an upper bound (here r=1.0), which will naturally reduce differences between predictive methods as they approach that upper bound. We next sought to test whether there was still a task-state FC advantage with larger amounts of data. A task-general approach was used in order to allow for more task data, but using all task data (including the to-be-predicted task) this time to include even more data. We varied the amount of rest and task-general data across 5, 10, 20, and 30 minutes of data. Both resting-state FC and task-general FC predictions accuracies increased significantly (all p<0.00001) with each incremental increase in data (**Figure 11**). An ANOVA indicated there was a main effect of amount of data (F(3)=9021, p<0.00001), a main effect of FC state (F(1)=1449, p<0.00001), and an interaction between amount of data and FC state (F(1,3)=46, p<0.00001). The interaction reflected the reduction in the task-general FC advantage relative to resting-state FC as the amount of data increased. Importantly, however, task-general FC predictions were significantly better at all amounts of data (all p<0.00001), including the highest amount of data (r=0.91 for task vs. r=0.86 for rest). These results demonstrate that the task-state advantage over resting-state FC for activity flow predictions remains regardless of the amount of data, though the task-state FC advantage naturally reduces at high amounts of data as the maximum possible prediction accuracy is approached.

**Figure 11.**
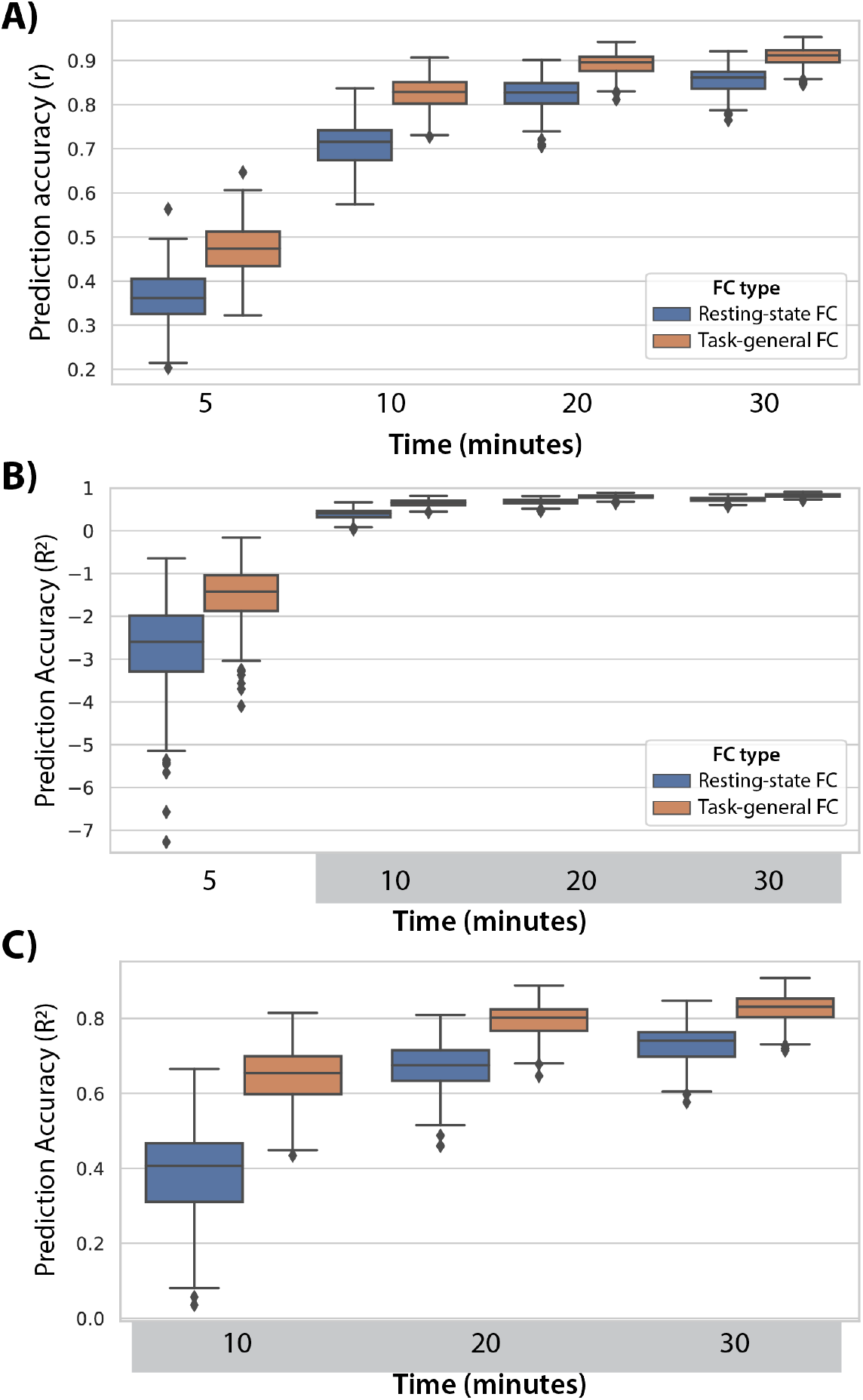
Activity flow prediction accuracy increases with the amount of data contributing to FC estimates. **A**) Prediction accuracies reported as Pearson correlations, collapsed across all 360 nodes and 24 task conditions. Task data was included from all 24 task conditions, similar to the prior task-general analyses but also including the to-be-predicted condition (to include more data). **B**) The same results reported as unscaled R^2^ (not r^2^), which can be interpreted (once multiplied by 100) as percent variance of the to-be-predicted activity pattern. These values range from negative infinity to 1, with the negative values quantifying how much worse the predictions are than predicting the mean of the data. **C**) The same results as in panel B, but with the 5 minute results excluded to improve legibility of the other results.

### Controlling for potential spatial smoothness confounds

It is thought that fMRI BOLD data have an inherent smoothness that partially reflects spatial properties of vasculature rather than neural activity (Lee et al., 1995; Menon and Kim, 1999). This smoothness is thought to be somewhere between 2 mm and 5 mm (Malonek and Grinvald, 1996; Logothetis and Wandell, 2004). The prior results might have been biased by this effect, given that the edge of each region is within 2 mm of the edge of nearby regions, introducing potential circularity in the prediction accuracies (Kriegeskorte et al., 2009; Cole et al., 2016).

We repeated the main analyses reported above with this bias removed. This was done by excluding all parcels with any vertices within 10 mm of each to-be-predicted region from the set of predictors (see Methods). We chose 10 mm to remain conservative, given uncertainty regarding the degree of vascular-driven smoothness of fMRI signal at any given location. Results were highly similar with this modification, despite including fewer predictors. Specifically, as hypothesized, we again found with multiple-regression FC that task-state FC improved activity flow-based predictions of task-evoked activations relative to resting-state FC. When using task-state FC: r=0.74, t(175)=139, p<0.00001. When using resting-state FC: r=0.39, t(175)=73, p<0.00001. The direct contrast between activity flow predictions with task-state vs. resting-state FC: r-difference=0.35, t(175)=99, p<0.00001. These results further confirm our primary hypothesis: That task-state FC is more informative regarding the paths of task-evoked activity flow than resting-state FC.

### Assessing prediction accuracy relative to repeat reliability (the noise ceiling)

A common approach for assessing the performance of predictive models is to compare predictions to the repeat reliability of the data, indicating how far the prediction accuracies are from the theoretical limit (the “noise ceiling”) (Naselaris et al., 2011; Nili et al., 2014). We reran the main analysis above (comparing predictions using multiple-regression FC with task vs. rest data), but building the activity flow models using the first task run and testing prediction accuracy based on task activations from the second task run. This was equivalent to a double cross-validation, with both the to-be-predicted region held out (as with standard activity flow mapping) as well as holding out the data used for estimating activations and FC of all regions. Another advantage of this approach was to better equate the resting-state FC estimates and task-state FC estimates, since (unlike task-state FC) resting-state FC was necessarily estimated in a separate fMRI run from the to-be-predicted task activations. Further, the prediction accuracies estimated in this manner are likely to generalize better to other independent data (in contrast to most of the prior predictions being optimized to the particular states subjects were in during task performance). Notably, since (1) error can come from the contributing activations in addition to the FC values, and (2) only half the data was used for both the FC and task activation estimates with this approach, we expect the predictions to be worse than most of the previous analyses. Note that the results in Figure 9 also used only a single task run per subject, but (unlike here) activation contributing to the activity flow mapping step still came from the same task run.

We started by calculating the repeat reliability between the two task runs across all nodes and task conditions (correlation between the actual task activations for run 1 and run 2). This was highly statistically significant, yet with an effect size lower than the activity flow-based predictions above: mean r=0.52, t(175)=65, p<0.00001. This r-value being far from r=1.0 suggests that reducing the amount of data by half likely reduces the reliability of the data substantially. Activity flow predictions with task-state FC were successful despite using half the data and predicting task activations in a different fMRI run: mean r=0.40, t(175)=57, p<0.00001. Importantly, the task-state FC boost to prediction (relative to resting-state FC) remained: mean r-difference=0.13, t(175)=38, p<0.00001. Condition-wise response profiles were also predicted better (each p<0.05, FWE corrected) with task-state FC than resting-state FC for 48% of the 360 cortical regions analyzed here: mean r-difference=0.09, t(175)=31, p<0.00001. These results demonstrate that the increase in prediction accuracy when using task-state FC versus resting-state FC was robust to predicting data in a separate task fMRI run.

## Discussion

The functional relevance of task-state FC has recently been called into question, based on the small size and number of task-related FC changes from resting-state FC (Cole et al., 2014; Krienen et al., 2014; Ito et al., 2020a) and the surprising efficacy of using resting-state FC to predict task activations (Cole et al., 2016; Tavor et al., 2016). Further, running counter to the common intuition that FC should increase as neural populations interact during task performance, FC tends to decrease as neural populations increase their activity levels and interact more strongly (Cohen and Maunsell, 2009; Ito et al., 2020a). We sought to test the possibility that – despite their counterintuitive nature – task-state FC changes nonetheless play an important role in shaping cognitive task activations. Supporting this conclusion, parameterizing network models using task-state FC consistently improved (relative to using resting-state FC) predictions of held-out cognitive task activations. Further, the use of empirically-derived network models provides mechanistic insight into how task-state FC influences brain function: by dynamically shifting the flow of activity between brain regions to shape task activations in a context-dependent manner.

This increase in prediction accuracy generalized across multiple tests, demonstrating the robustness of this effect. First, cognitive task activations across all 24 task conditions and 360 cortical brain regions tested were better predicted when using Pearson-correlation FC (the field standard) estimated during task performance relative to resting state. Importantly, mean task-evoked activations (i.e., the to-be-predicted signals) were aggressively removed prior to task-state FC estimation, eliminating analysis circularity and leaving only spontaneous and induced activity to contribute to task-state FC estimates (Cole et al., 2019). Despite using such “background” connectivity (also termed “noise correlations”) (Norman-Haignere et al., 2012) the task-state FC estimates nonetheless better described task activity flow than resting-state FC. Notably, the vast majority of studies that have suggested high functional relevance of task-state FC due to its correlation with individual differences in behavior (e.g., Greene et al., 2018) did not remove task activation variance (already known to be highly related to behavior), such that these previous findings did not conclusively demonstrate functional relevance of task-state FC. Second, using multiple-regression FC to reduce causal confounds (Reid et al., 2019; Sanchez-Romero and Cole, 2020) further improved predictions of cognitive task activations, suggesting (tentatively for now) the causal plausibility of activity flow predictions using task-state FC. Third, task-state FC from tasks other than the to-be-predicted task also provided a boost in activation prediction accuracy, though it did not provide as strong a boost in accuracy as when the same task was used.

The present results are consistent with a variety of studies showing that – despite their small size and number – task-related FC changes nonetheless reliably differ from resting-state FC (Cole et al., 2014; Krienen et al., 2014; Gratton et al., 2018). This reliability across time and also (to some extent) across subjects suggests there may be an important functional role for task-state FC. Importantly, we also found that individual differences play a large role in the task-state FC boost in activation prediction accuracy. This is consistent with results showing that task-state FC changes are largely subject-specific (Gratton et al., 2018). This suggests that group-averaged task-state FC results will always partially mischaracterize task-state FC changes, and that individualized characterization (and/or characterization of individual variation) will be especially important in future work. Note that (unless noted otherwise) the activity flow mapping results were based on individualized FC and activations, with predicted-to-actual comparisons made prior to group averaging.

The present results advance our understanding of task-state FC by mechanistically simulating how these estimated connections may support task-evoked activations. This is an advance because of theoretical and mechanistic uncertainties regarding task-state FC, which contrasts with theoretical and mechanistic certainties regarding task-evoked activations. For instance, activations are known to cause perception, action, and cognition (e.g., via transcranial magnetic stimulation or intracranial stimulation) (Valero-Cabré et al., 2017). There is less mechanistic certainty regarding fMRI-based activations relative to neural spike activations (i.e., increases in spike rate), yet it is well established that neural spike activations can cause fMRI activations (Logothetis et al., 2001; Lee et al., 2010). Supporting this interpretation of fMRI activations, many highly replicated fMRI activations are consistent with extremely well-established causal mechanisms (based on over a century of evidence from more invasive approaches), such as activations in primary motor cortex during motor action (Yokoi et al., 2018), primary visual cortex during visual perception (Wandell and Winawer, 2015), and primary auditory cortex during auditory perception (Striem-Amit et al., 2011). Yet fMRI is known to be sensitive to other factors besides spiking activity, such that spikes cannot be inferred with certainty based on fMRI observations. Including nuisance regression to remove other signals that fMRI is sensitive to (e.g., physiological artifacts) – as we have here – can help with this kind of inference, however.

Relative to activations, little is known regarding the mechanisms underlying FC, such as the task-related changes in correlations/regressions investigated here. In the non-human animal literature task-state FC is typically termed “noise correlations”, because they are calculated (as we have here) as covariance above and beyond the “signal”: cross-trial mean task-evoked activations. It is well known that noise correlations between neurons tend to decrease from rest to task (Cohen and Kohn, 2011), and that this tends to increase the information capacity of neural populations via making neural responses more distinct from each other (Averbeck et al., 2006; Cohen and Kohn, 2011). We recently showed that such task-related decreases in correlation generalize to correlations between brain regions (not just individual neurons) in both monkey spiking data and human fMRI data (Ito et al., 2020a) (**Figure 2**). Beyond this abstract information theoretic interpretation, however, it has been unclear whether task-state functional connections (noise correlations) actually contribute to cognitive/perceptual/motor functionality. Here we provided evidence consistent with this possibility, given that task-state FC better predicts task-evoked activations (which are more directly and mechanistically related to cognitive/perceptual/motor functionality) than intrinsic FC.

We sought to improve theoretical insights into the mechanistic role of task-state FC using the concept of activity flow. Activity flow (as a theoretical construct) is the movement of activity amplitudes between neural populations, which activity flow mapping attempts to quantify. Since true activity flow involves directionality and spatiotemporal precision not currently possible with fMRI, we previously validated its use with fMRI data using neural mass simulations. We found that simulated fMRI data, while imperfect, was quite accurate at predicting ground-truth activity flows in the simulated data (Cole et al., 2016; Ito et al., 2017). Standard neuroscience theory attributes the flow of neural activity solely to action potentials flowing over axons, which cause the release of neurotransmitters at synapses (Hodgkin and Huxley, 1952). These neurotransmitters are then thought to cause the BOLD signal via interactions with nearby astrocytes and blood vessels (Attwell et al., 2010). Thus, standard neuroscience theory provides a straightforward mechanistic account of how activity flow relates to fMRI BOLD: local activity changes – measured indirectly via fMRI BOLD – can be related to each other via activity flows over long-distance connections (action potentials over axons). However, fMRI BOLD is susceptible to artifacts and distortions due to spatiotemporal downsampling and other factors. We used rigorously validated data preprocessing to reduce the impact of artifacts, while we relied on well-established theory and the validation simulations mentioned above (which also involved spatiotemporal downsampling to simulate fMRI) to support our fMRI-based inferences. Notably, many of the conclusions reached here were dependent on the extensive spatial coverage (the entire neocortex), relatively high spatial localization certainty (relative to, e.g., electroencephalography), along with moderately high temporal resolution (seconds not minutes) afforded by fMRI.

Conclusions must nonetheless be restricted to some extent given limitations of fMRI BOLD relative to ground-truth spiking activity. For instance, the FC measures used do not estimate causal directionality, such that the directions of estimated activity flows were unknown. Notably, however, most cortico-cortical connections are known to be bidirectional (Markov et al., 2014). This suggests that most of the activity flow predictions are at least partially correct under the assumption of bidirectionality. Further, since resting-state FC (the baseline for comparison) also had the directionality limitation, our conclusions are likely unaffected by the lack of directional information. Another limitation of fMRI BOLD is that it has an inherent smoothness that partially reflects spatial properties of vasculature rather than neural activity (Lee et al., 1995; Menon and Kim, 1999). This effect is thought to be between 2 mm and 5 mm (Malonek and Grinvald, 1996; Logothetis and Wandell, 2004). We ruled this out as a major factor here by conducting a follow-up analysis in which we removed all parcels within 10 mm of the to-be-predicted activations from the activity flow mapping procedure.

Building on our recent non-human (and human) primate study showing that most correlation-based functional connections decrease from rest to task (Ito et al., 2020a), we found that most regression-based functional connections also decrease. These results appear to run counter to the common intuition that tasks should increase neural interactions overall. Activity flow mapping adds insight here, however, since we found that activity flow (*activations* ✕ *FC*) often increases even as FC decreases (see **Figure 1C & 1D**). Thus, the common intuition that neural interactions (quantified as activity flows) often increase during tasks relative to rest can stand, while acknowledging that most functional connections (correlations and regressions) decrease from rest to task states. Notably, our recent study (Ito et al., 2020a) suggested that these task-state FC decreases likely result from neural populations being related via sigmoidal transfer functions, suggesting the importance of future work examining estimation of task-state FC using nonlinear regression approaches. Even if a single nonlinear function could account for both resting-state FC and task-state FC, however, the insights gained via changes in separate piece-wise linear estimates (as done here; see Figure 2C) would remain valid.

In conclusion, we found strong evidence consistent with task-state FC having a prominent role in neurocognitive functions. This built on our prior work suggesting intrinsic brain connectivity (as measured by resting-state FC) has a large role in neurocognitive functionality, by shaping cognitive task activations (Cole et al., 2016; Ito et al., 2017; Mill et al., 2019). We again quantified the likely contribution of connectivity to cognitive task activations, finding that task-state FC consistently improved prediction of task activations relative to resting-state FC. This suggests that task-state FC likely has an important functional role despite changes from resting state being small in size (Cole et al., 2014; Krienen et al., 2014), and despite most of those (statistically significant) changes being reductions in FC strength. This conclusion is highly general, as cognitive task activations were better predicted across all 360 cortical regions and all 24 task conditions tested. We anticipate these findings to facilitate future interpretation of task-state FC effects, such as the interpretation of task-state FC decreases in the context of increased activity as potentially reflecting overall increases in activity flow.

## Acknowledgements

The authors thank Ethan McCormick for helpful feedback. The authors acknowledge support by the US National Institutes of Health under awards R01 AG055556 and R01 MH109520. Data were provided, in part, by the Human Connectome Project, WU-Minn Con-sortium (Principal Investigators: D. Van Essen and K. Ugurbil; 1U54MH091657) funded by the 16 NIH Institutes and Centers that support the NIH Blueprint for Neuroscience Research; and by the McDonnell Center for Systems Neuroscience at Washington University. The authors also acknowledge the Office of Advanced Research Computing (OARC) at Rutgers, The State University of New Jersey for providing access to the Amarel cluster and associated research computing resources that have contributed to the results reported here. The content is solely the responsibility of the authors and does not necessarily represent the official views of any of the funding agencies.

## Notes

### Competing Interest Statement

The authors have declared no competing interest.

